# A Benchmark of Unsupervised Circadian Phase Inference Methods for Bulk Transcriptomic Data

**DOI:** 10.64898/2026.10.08.757747

**Authors:** Souparna Purohit, Gregory R. Grant, Thomas G. Brooks

## Abstract

Circadian phase can be inferred from gene expression when sample collection times are unavailable, but the available methods have not been systematically compared. We compared ten unsupervised phase inference methods and two clock-gene baselines on 25 datasets with reference times (16 real datasets from five studies and 9 simulations). The primary measure was the aligned circular mean absolute error (MAE), with a permutation test for significance, circular correlations and gene-level reconstruction of rhythmicity, acrophase and amplitude were secondary outcomes. CHIRAL had the lowest mean MAE (1.80 h, 95% confidence interval 1.21-2.97 h from resampling study families). The two clock-gene baselines (2.15 and 2.20 h) were next, followed by TEMPO, CYCLOPS and CYCLOPS 2.0 (2.49-2.80 h), and the confidence intervals of these six overlapped substantially. The other six methods had mean errors of 3.91-4.86 h, compared with 5.08-5.23 h for permuted phases. Accuracy depended on the dataset and on the strength of clock-gene rhythms, and, for several methods, on the random seed and the adjustment for batch or covariates. When the inferred phases were used to analyse gene rhythms in 22 datasets, the four leading methods identified rhythmic genes with a median ROC AUC of 0.64-0.70 and recovered gene acrophases with a median error of 0.18-0.64 h. In two mouse datasets without collection times, agreement with external reference acrophases differed between methods and tissues.

## Introduction

Gene expression exhibits approximately 24-hour rhythms, with rhythmic genes and their peak expression times varying across tissues. These rhythms include genes encoding drug targets and drug-metabolizing enzymes, with implications for the timing of treatment (Ruben et al. 2018). Characterizing these rhythms across samples requires information about collection time or an estimate of each sample’s circadian phase.

Collection times are often unavailable in public transcriptomic datasets, including biopsy collections and clinical or population cohorts. Some post-mortem studies record the time of death, which can provide a temporal reference for expression analyses (Ketchesin et al. 2023). Moreover, collection and death times are reference times for a sample, not a measurement of the donor’s internal circadian phase.

When these sample circadian times are not known, temporal information may be inferred from the data, and has been used to reconstruct gene rhythms (Anafi et al. 2017). Critically, these methods have enabled the analysis of circadian rhythms relevant for human health in tissues or conditions that cannot be collected in controlled experimental settings due to ethical or practical considerations. For example, inferred samples times have found sex- and age-specific trends in rhythms (Talamanca et al. 2023), altered rhythms in luminal A breast cancers cells (Li et al. 2024), and the effects of Alzheimer’s disease on human brain cells (Hollis et al. 2025).

Methods for estimating circadian phase from gene expression include supervised prediction and unsupervised phase reconstruction. Supervised methods use labelled reference data to learn an expression-time relationship and apply the fitted model to new samples; examples are the molecular timetable (Ueda et al. 2004), ZeitZeiger (Hughey et al. 2016) and TimeSignature (Braun et al. 2018). Their applicability depends on how well that relationship transfers to the tissue, species and study being analysed. Unsupervised methods estimate sample phases from an expression matrix without using the samples’ collection times, often with prior information about clock genes or rhythmic expression. Unsupervised reconstruction is particularly useful when collection times and a suitable labelled training dataset are unavailable. Methods of this kind include CYCLOPS (Anafi et al. 2017), CYCLOPS 2.0 (Li et al. 2024), CHIRAL (Talamanca et al. 2023), TEMPO (Auerbach et al. 2022), COFE (Ananthasubramaniam and Venkataramanan 2025), DCPR (Han et al. 2026), PENN (Ansary Ogholbake and Cheng 2023, 2024) and PROTECT (Ansary Ogholbake and Cheng 2025). We evaluated PENN using the released implementation of the 2023 conference paper (Ansary Ogholbake and Cheng 2023); the code of the 2024 journal version (Ansary Ogholbake and Cheng 2024) was not released. Cyclum (Liang et al. 2020) and reCAT (Liu et al. 2017) were developed for the cell cycle in single cells and address a related circular-ordering problem.

The authors of these methods evaluated them on their own datasets, with their own preprocessing and varying accuracy measures. These papers have also compared their methods to selected phase-inference approaches (Han et al. 2026; Auerbach et al. 2022), but differ in the methods, datasets, preprocessing and evaluation conventions used. A comparable evaluation exists for the related problem of detecting rhythmic genes when times are known (Mei et al. 2021). However, evaluations performed by methods developers upon their own methods have been shown to be biased (Buchka et al. 2021), and there has been no systematic, independent evaluation of these methods to date.

Inferred phases are often used to estimate gene rhythmicity, amplitude and acrophase in datasets without temporal annotations. Errors in sample phase can therefore affect downstream conclusions even when the inferred ordering appears plausible. Evaluating both sample phase accuracy and gene-level rhythm reconstruction is necessary to assess the usefulness of these estimates.

Here we compare ten unsupervised phase inference methods and two simple clock-gene baseline methods on 25 datasets with reference times, using a common evaluation protocol across real and simulated datasets. We used aligned circular mean absolute error as the primary performance measure, with circular correlations and gene-level rhythm reconstruction as secondary outcomes. A permutation analysis provided a dataset- and method-specific reference for performance relative to chance. We ask how accurate the methods are relative to chance and to the baselines, whether gene-level rhythms are reconstructed, how accuracy depends on the dataset, and how sensitive the results are to the random seed, the seed gene list and batch or covariate adjustment. We used simulations to determine sensitivity to parameters such as rhythm shape which cannot be controlled in real datasets (Sandve et al. 2022). We also evaluated the methods against reference acrophases in two expression datasets from Diversity Outbred mice, comprising liver and adipose tissue, that have no collection times.

## Results

### Benchmark design

The main comparison included 25 datasets: 16 real datasets from five studies and 9 simulations (Fig. 1a,b). These were grouped into 6 families of related datasets (such as different tissues from the same study). The real datasets were human and mouse tissues with recorded sampling or death times, referred to here as reference times (Fig. 1b). Two expression datasets from Diversity Outbred (DO) mice, comprising the liver and adipose tissues of 466 mice, have no collection times. They were evaluated against external reference acrophases and amplitudes. The datasets differed widely in how strongly twelve canonical clock genes cycle against reference time (median partial R^2^ from 0.08 to 0.91; Fig. 1b).

**Figure 1.**
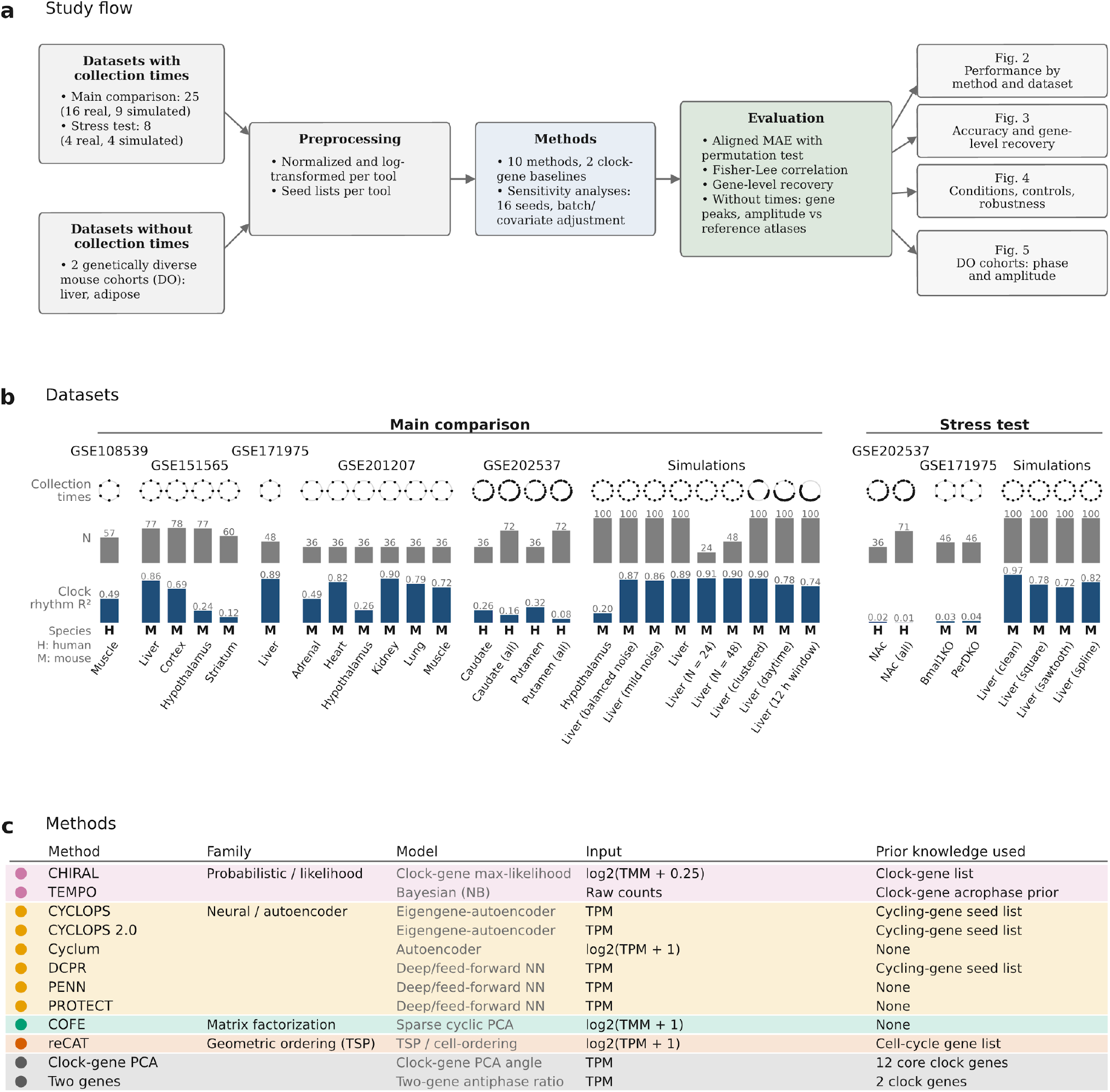
Study design. (a) Datasets, preprocessing, methods and evaluation, and the figures that report each analysis. (b) Datasets with collection times, in the main comparison and the stress test: collection times around the 24 h cycle (rings, one dot per sample), number of samples (N), clock-gene rhythm strength (median cosinor R^2^ of twelve clock genes against collection time) and species. (c) The methods and the two clock-gene baselines, with model family, model, input matrix and prior knowledge used.

We evaluated ten methods and two baselines (Fig. 1c). The methods are probabilistic (CHIRAL, TEMPO), neural-network or autoencoder based (CYCLOPS, CYCLOPS 2.0, Cyclum, DCPR, PENN, PROTECT), matrix factorization (COFE) or geometric ordering (reCAT). Methods were run as described in their publications (Supplementary Methods). The two baseline methods assigned each sample the angle of its position in the plane of the first two principal-component scores of twelve standardized clock genes (clock-gene PCA), or in the plane of the standardized expression of ARNTL and NR1D1 (two genes).

To verify that methods were being used as intended by their original authors, we attempted to replicate published results where these were available (Supplementary Table S1). We succeeded in producing similar results in 8 methods, but we were not able to reproduce the results for PENN and PROTECT (Supplementary Table S1).

Inferred phases were scored, for each dataset and method, using the aligned circular mean absolute error (MAE): the mean shortest circular distance, in hours, between inferred phases and reference times after optimizing a global rotation and reflection (Fig. S1). We permuted each run’s inferred phases and repeated the alignment to obtain its null distribution, yielding a p-value from a one-sided test.

### Accuracy compared with chance and with clock-gene baselines

The primary ranking used the mean aligned MAE across the 25 datasets, each weighted equally, with confidence intervals obtained from resampling whole study families to allow for dependence within studies (Fig. 2a, medians and other measures in Supplementary Table S2). The methods fell into three groups.

**Figure 2.**
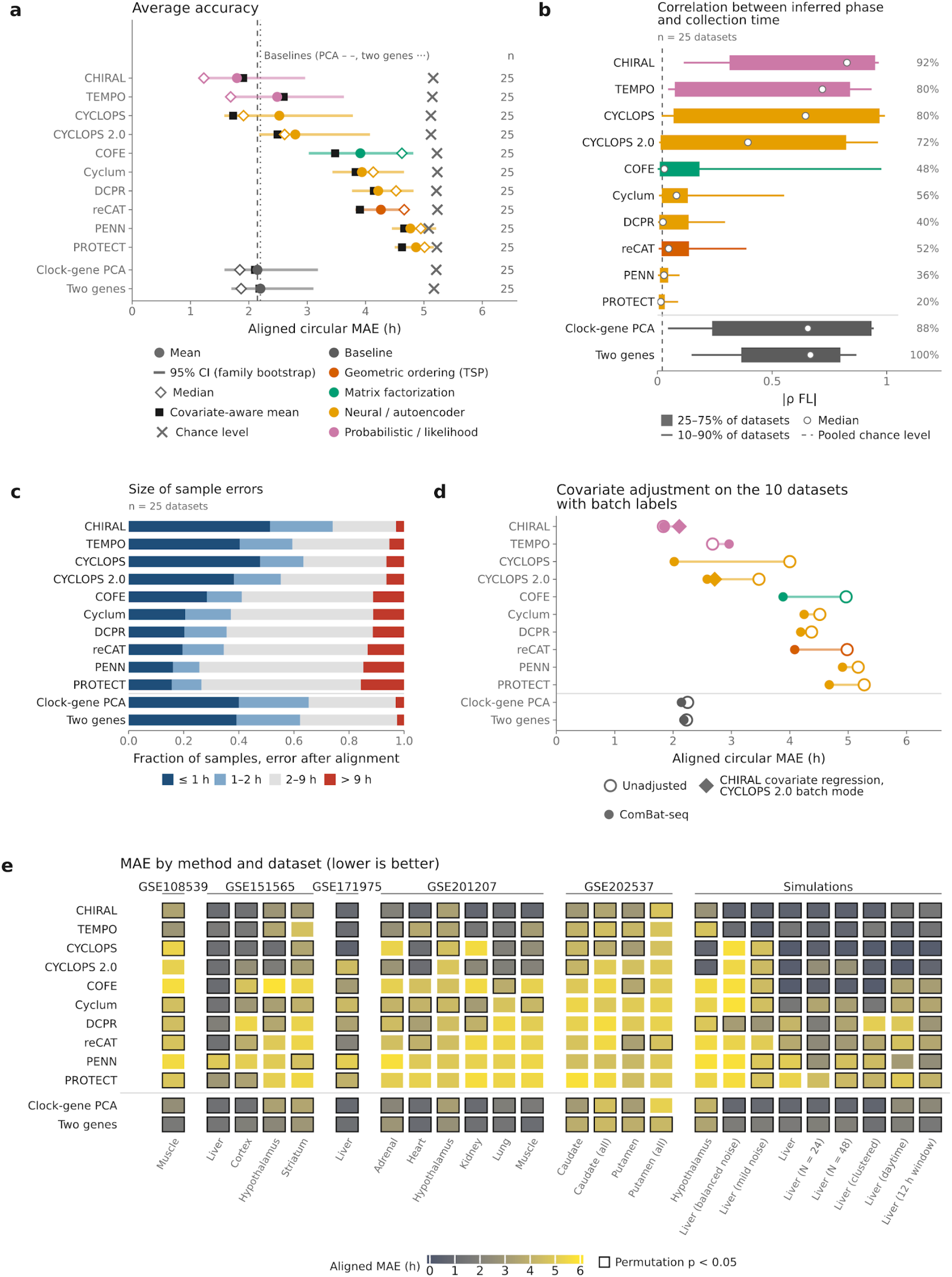
Overall performance in the main comparison. (a) Left: Mean aligned MAE per method (dot) with the 95% family-bootstrap interval (horizontal line), median (open diamond), mean with covariate-adjusted results substituted where available (black square) and chance level (cross: the mean aligned MAE after shuffling the method’s inferred phases among samples, averaged over datasets). (b) Absolute Fisher-Lee correlation between inferred phases and collection times over the main-comparison datasets. Percentages give the share of datasets in which the correlation passed the run-specific permutation criterion (at permutation p < 0.05). (c) Fraction of samples in each aligned-error band, mean over datasets. (d) Mean aligned MAE on the ten datasets with batch labels: unadjusted (open circle), on ComBat-seq-corrected input (filled circle) and, as diamonds, CHIRAL with covariate regression and CYCLOPS 2.0 in batch mode. (e) Aligned MAE of each method on each dataset, grouped by study family. Dark outline indicates permutation p < 0.05.

CHIRAL had the lowest error (1.80 h; 95% confidence interval 1.21-2.97 h) and was the only method ahead of both clock-gene baselines (2.15 and 2.20 h). TEMPO, CYCLOPS and CYCLOPS 2.0 followed at 2.49-2.80 h, behind the baselines on the mean. Based on the median, TEMPO (1.69 h) moved ahead of the baselines while CYCLOPS (1.91 h) was level with them. The remaining six methods’ mean errors of 3.91-4.86 h were not far below their chance level of 5.08-5.23 h.

### Sample-level accuracy

The absolute Fisher–Lee correlation between inferred phases and collection times ranked the methods much as the MAE did. In particular, it also splits the methods into two groups (Fig. 2b). The top group consists of CHIRAL (0.83 median), TEMPO (0.72), CYCLOPS (0.65), and CYCLOPS 2.0 (0.39), while the bottom group had the rest of the tools, all at 0.08 or lower. The two baselines were at 0.66 and 0.67. Another metric we measured is the fraction of samples within 1h of reference time (after optimal alignment), averaged across datasets. The two groups above stayed consistent: CHIRAL (51%), CYCLOPS (48%), TEMPO (40%), and CYCLOPS 2.0 (38%) were again the top performers, followed by the rest of the tools at 16-28% (with COFE at 28%). Meanwhile, the two baselines were at 40% (clock gene) and 39% (two-gene) (Fig. 2c). The mean fraction of samples with errors above 9 h, which places a sample near the opposite side of the cycle, was 3-6% for CHIRAL, TEMPO, CYCLOPS, CYCLOPS 2.0 and the baselines and 11-16% for the other six methods.

The ordering within these groups is uncertain. The confidence intervals overlap substantially, although CHIRAL had the lowest mean in 92% of 5,000 resamples of study families. A Friedman permutation test across the six study families indicated that the methods differed (p < 0.001) - that is, the ordering of the methods was consistent across families. In post-hoc Nemenyi comparisons, however, only contrasts between the top and the bottom of the ranking reached p < 0.05: CHIRAL against COFE, PENN and PROTECT (p = 0.022, 0.001 and 0.003), and each baseline and CYCLOPS against PENN (p = 0.022 and 0.037), with the baselines also differing from PROTECT (p = 0.037). Note that the absence of a detected difference does not establish that two methods perform comparably.

Agreement with collection times was often statistically detectable even when the error was large. In the one-sided aligned-MAE permutation test (p < 0.05, unadjusted), CHIRAL was significant on all 25 datasets and TEMPO, CYCLOPS and CYCLOPS 2.0 on 19-23, but the six lower-ranked methods were also significant on 9-18 datasets while their mean errors were only 0.3-1.3 h below chance (Fig. 2a).

### Batch and covariate correction

The main comparison gave the methods no batch or covariate annotations. Ten datasets had a batch-like variable (age group in the six GSE201207 tissues, disease state in the two GSE202537 regions and the simulated batch in the two simulated datasets), and on these we compared each method’s unadjusted run with its run on ComBat-seq-corrected counts. CHIRAL was also run after regressing out the batch (and an additional post-mortem interval covariate for the two GSE202537 tissues) following the authors’ workflow, and CYCLOPS 2.0 was also run with the batch covariate as well as the post-mortem interval for GSE202537) (Fig. 2d; per-dataset results in Supplementary Fig. S12; Methods Table 5). Adjustment helped some methods substantially and others not at all. ComBat-seq halved the mean error of CYCLOPS on these datasets (4.00 to 2.02 h, lower on 8 of 10) and lowered that of reCAT, COFE and CYCLOPS 2.0 by 0.9-1.1 h, but left CHIRAL unchanged (1.83 and 1.84 h) and raised the error of TEMPO (2.68 to 2.96 h). Neither built-in adjustment outperformed ComBat-seq: CHIRAL’s covariate regression increased its error to 2.11 h, and CYCLOPS 2.0’s batch mode gave 2.71 h against 2.58 h with ComBat-seq. Substituting adjusted results on these ten datasets into the 25-dataset means (black squares in Fig. 2a) moves CYCLOPS from 2.52 to 1.73 h, below CHIRAL at 1.90 h, but the family-bootstrap intervals overlap widely (0.77-3.42 and 1.23-3.29 h), and because the substituted adjustment differs between methods this is not a ranking under one common procedure.

### Variability by dataset

Accuracy also varied widely across datasets (Fig. 2e). At least one method had an error near or below 1 h on mouse liver, heart, kidney and muscle datasets, whereas on the four human caudate and putamen datasets (GSE202537) no method or baseline had an error below 2.9 h. These datasets have among the weakest clock-gene rhythms in the benchmark (Fig. 1b, Fig. S7), which is consistent with the higher error.

Their reference times are also less precise (times of death, known to within about 2 h for unwitnessed deaths), although this uncertainty is too small to explain errors of 3-4 h. More generally, the absolute performance on datasets tracked with the strength of clock-gene rhythms within a dataset (Fig. S6c and Fig. S7): datasets with weaker rhythms had higher error (Spearman ρ = -0.58 between rhythm strength and the median error over methods). In addition, errors were about twice as large on real as on simulated datasets for the leading methods (Supplementary Table S2).

Further tests of variability of the tool rankings are performed by leave one out tests, as well as ranking the tools by their average MAE on various families of datasets, such as tissues or organisms (Fig. S6). Overall, despite the variability, CHIRAL was the most consistent, ranking among the four best methods in every dataset, whereas every other method ranked seventh or lower in at least one. In particular, CHIRAL had the lowest error in 11 of the 16 real datasets. On the simulations, where the leading methods were all within about an hour of one another, CYCLOPS was usually lowest (6 of 9). It also remained first by mean error when any one study family was left out and on the real datasets alone (Fig. S6).

### Rhythm reconstruction

Next we asked how well the inferred phases could reconstruct gene-level rhythms. For each dataset, reference rhythmicity, acrophase and amplitude were taken from cosinor fits to the collection times, and the same fits were repeated using each method’s inferred phases (Supplementary Methods). Fig. 3a shows the clock genes for the best and worst methods on the dataset where the two differ most (GSE201207 mouse kidney).

**Figure 3.**
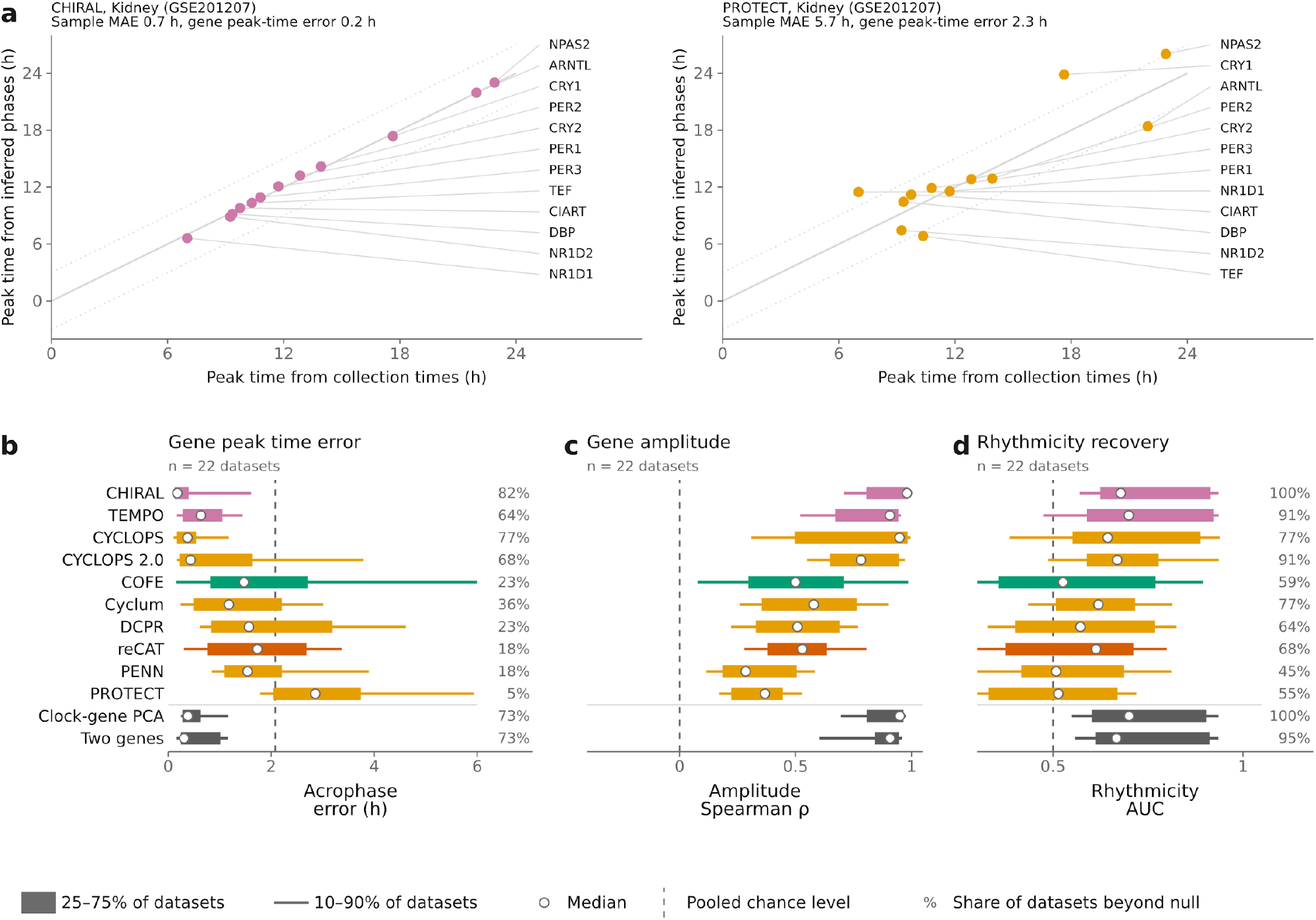
Sample-level accuracy and gene-level rhythm reconstruction. (a) Example of clock-gene peak times fitted to the inferred phases against those fitted to the collection times (aligned by rotation and reflection), for the best and worst method by MAE on one dataset (GSE201207 kidney). (b–d) Gene-level reconstruction result, which compare the gene rhythms obtained from inferred-phase fit to fits using reference phases: median acrophase error (b), amplitude Spearman correlation (c) on reference-rhythmic genes, and rhythmicity ROC AUC (d) (reference-rhythmic genes that fail the acrophase standard-error criterion are omitted from the AUC; Supplementary Methods). In (b–d) dots are medians over datasets (22), bars the 25th–75th and lines the 10th–90th percentile. Dashed lines are chance levels: the permutation null averaged over methods and datasets in (b), 0 in (c) and 0.5 in (d). Percentages give the share of datasets in which the metric passed the run-specific permutation criterion (acrophase error below the 5th percentile of the null, AUC above the 95th percentile).

Gene peak times separated the methods in the same way as sample-level accuracy. Across the 22 datasets with reference-rhythmic genes, the median acrophase error was 0.18-0.64 h for CHIRAL, CYCLOPS, CYCLOPS 2.0 and TEMPO, and these methods fell below the 5th percentile of their own permutation null in 64-82% of datasets (Fig. 3b). For the other six methods the median error was 1.18-2.86 h, against a pooled permutation reference of about 2.1 h, and they passed the same criterion in at most 36% of datasets. The two clock-gene baselines matched the leading methods (0.31 and 0.38 h, on 73% of datasets).

The median correlation between reconstructed and reference amplitudes was 0.78-0.98 for the four leading methods and the baselines, and remained positive for the others (0.28-0.58), including methods whose peak-time errors were near chance (Fig. 3c). Rhythmicity AUC discriminated least: medians ranged only from 0.51 to 0.70, although the AUC exceeded the 95th percentile of its null in all datasets for CHIRAL and in 45% for PENN (Fig. 3d).

Rhythm reconstruction results varied slightly under differing definitions of reference rhythmic genes (Fig. S5).

### Sensitivity to simulation parameters

We varied the sampling design, sample size and waveform one at a time relative to a reference liver simulation, which had uniform sampling, N = 100, cosine waveform (see Methods for details). Each condition was simulated once, so differences between conditions are not accompanied by an estimate of their uncertainty (Fig. 4a-c). On the reference simulation, CHIRAL, CYCLOPS, CYCLOPS 2.0 and COFE had errors of 0.3-0.6 h, TEMPO and Cyclum 1.5 h, and DCPR, reCAT, PENN and PROTECT 3.7-5.3 h.

**Figure 4.**
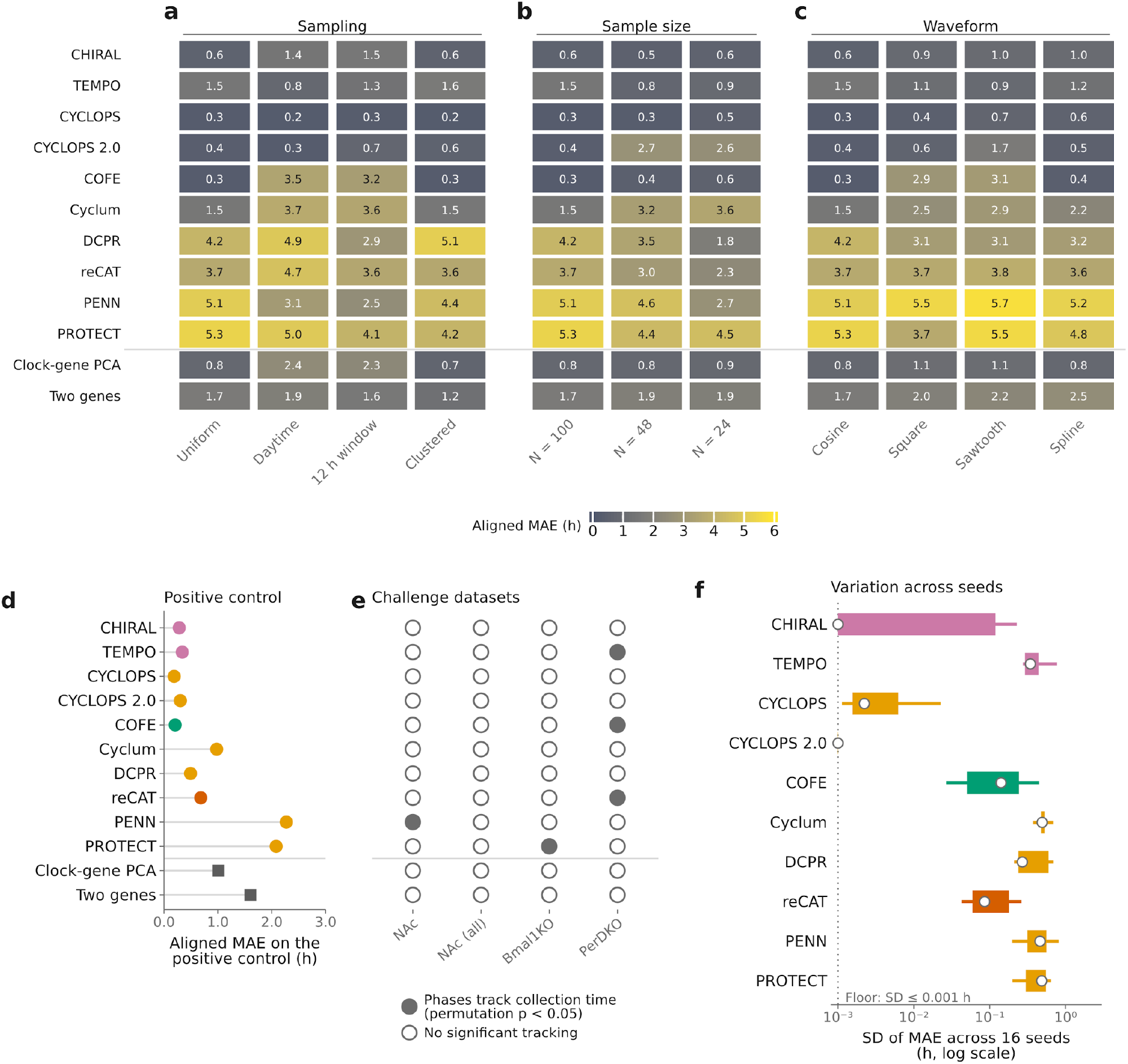
Simulation conditions, controls and sensitivity analyses. (a–c) Aligned MAE on simulated liver datasets that vary the sampling design (a), sample size (b) and waveform (c) from a reference simulation. (d) Aligned MAE on the positive-control simulation. (e) Agreement with collection times on the four challenge datasets (filled denotes significance in the one-sided aligned-MAE permutation test at p < 0.05). Standard deviation of aligned MAE across 16 random seeds on the seven repeat-run datasets (log scale, with ranges below 0.001 h drawn at the dotted floor line). Small ranges indicate reproducibility across seeds.

Restricting the sampling times affected some of the accurate methods strongly and others not at all. Sampling only during daytime or within a 12 h window raised the error of COFE from 0.3 h to 3.5 and 3.2 h, and roughly doubled or tripled that of CHIRAL and Cyclum, whereas CYCLOPS stayed at 0.2–0.3 h. Clustering samples around a few times changed the error of the accurate methods by at most 0.2 h.

Reducing the sample size from 100 to 48 and 24 mainly affected CYCLOPS 2.0 (0.4 h to 2.7 and 2.6 h) and Cyclum (1.5 h to 3.3 and 3.6 h); CHIRAL, CYCLOPS and COFE remained at about 0.6 h or below. DCPR and PENN had lower errors at N = 24 than at N = 100 (4.2 to 1.8 h and 5.1 to 2.7 h).

Non-sinusoidal waveforms were handled unevenly. Square and sawtooth waveforms raised the error of COFE to 2.9 and 3.1 h, and the sawtooth raised that of CYCLOPS 2.0 to 1.7 h, while CHIRAL, CYCLOPS and TEMPO stayed at about 1 h or below under all three alternative waveforms.

### Positive and negative control datasets

Next we created a positive-control simulation that has strong clock-gene rhythms and reduced count noise, representing a best-case scenario (see Supplemental Methods for details). Eight of the ten methods had an aligned MAE between 0.19 and 0.98 h, compared with 1.01 h for clock-gene PCA and 1.60 h for the two-gene baseline (Fig. 4d). PENN (2.27 h) and PROTECT (2.08 h) had higher errors than both baselines.

As a negative control, we used three “challenge” datasets in which clock-gene rhythms are weak or disrupted: human nucleus accumbens from GSE202537, analysed for control subjects alone (36 samples) and with psychosis subjects included (71 samples), and mouse liver from Bmal1-knockout and Per1/Per2 double-knockout animals (GSE171975, 46 samples each). No method showed significant agreement with the collection times on more than one dataset, and only five of 48 comparisons met the permutation p < 0.05 criterion (Fig. 4e).

### Sensitivity to seeds

To assess how much results vary with the random seed, we reran each method with 16 random seeds on seven datasets (four real and three simulated), while the main comparison (from Fig. 2 and Fig. 3) used just one seed. For each method and dataset we calculated the standard deviation of the aligned MAE across seeds (Fig. 4g). CYCLOPS 2.0 fixes its random seed internally, so its 16 runs were identical and its sensitivity to initialization could not be assessed. The median standard deviation over datasets was below 0.001 h for CHIRAL and 0.002 h for CYCLOPS, 0.09 h for reCAT and 0.14 h for COFE, and between 0.27 and 0.49 h for DCPR, TEMPO, PENN, PROTECT and Cyclum. CHIRAL was not uniformly stable: its standard deviation was below 0.001 h on four datasets but 0.28 h on one, where its error ranged over 0.68 h across seeds.

### Datasets without known collection times

For the two Diversity Outbred mouse datasets (GSE266549, GSE266569), we have no reference collection times. We compared the gene acrophases fitted to each method’s phases with reference acrophases, using the Zhang mouse tissue atlas (Zhang et al. 2014) for adipose and liver and, for liver, a separate liver reference assembled from the mouse liver atlas (Brooks et al. 2023) with the Zhang liver study excluded (leaving 1,088 samples from 56 studies; Fig. 5a–c).

**Figure 5.**
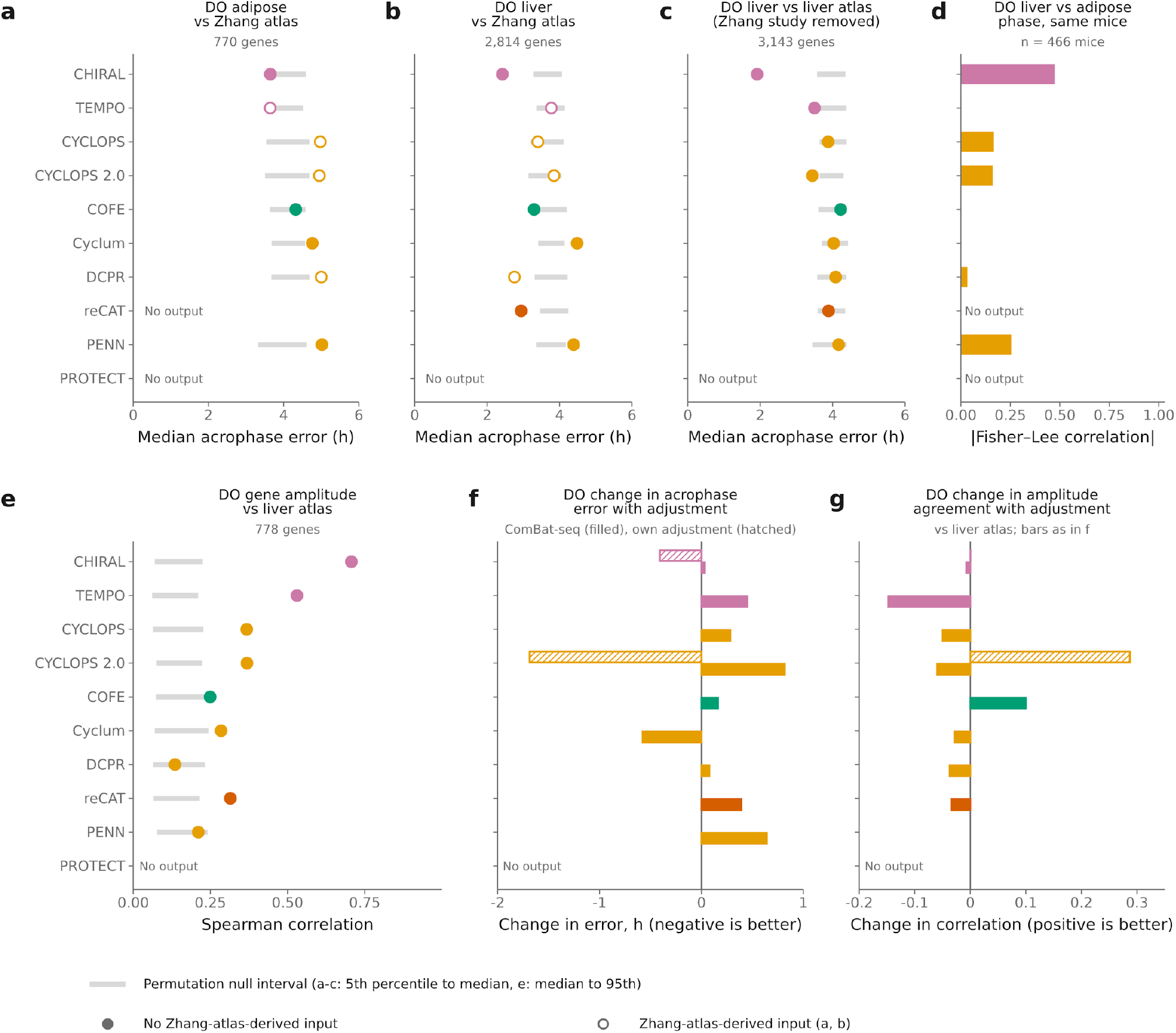
Agreement with external references in datasets without collection times. (a–c) Median acrophase error against reference acrophases for DO adipose and liver against the Zhang atlas (a, b) and DO liver against the separate mouse liver reference with the Zhang study removed (c) (grey bars are permutation null from 5th percentile to median), open circles are methods with Zhang-atlas-derived input. (d) |Fisher-Lee circular correlation| between inferred liver and adipose phases across the 466 mice. (e) Spearman correlation of gene amplitudes with the liver atlas (grey bars are permutation null from median to 95th percentile). (f) Change in the median acrophase error of (c) with adjustment (ComBat-seq adjustments are filled, and covariate regression for CHIRAL or batch mode for CYCLOPS 2.0 are hatched). (g) Change in the amplitude correlation of (e) with the same adjustments (bars as in f). Changes in (f, g) are relative to each method’s unadjusted run.

In adipose (770 rhythmic reference genes), no method’s median acrophase error fell below the 5th percentile of its permutation distribution (eight methods with output; Fig. 5a). In liver, CHIRAL, COFE, DCPR and reCAT placed below the 5th percentile of the permutation distribution in the Zhang atlas (2,814 rhythmic genes; Fig. 5b), as did CHIRAL, TEMPO and CYCLOPS 2.0 against the separate liver reference (3,143 rhythmic genes; Fig. 5c).

We also assessed whether phases inferred separately from liver and adipose expression were associated across the 466 mice profiled in both tissues. The absolute Fisher–Lee circular correlation between the two tissues’ phases was 0.47 for CHIRAL, 0.26 for PENN, 0.17 for CYCLOPS and 0.16 for CYCLOPS 2.0, and at most 0.03 for COFE, Cyclum, DCPR and TEMPO (Fig. 5d).

Reconstruction of gene amplitudes was best for CHIRAL (0.71 Spearman correlation to reference amplitudes) and TEMPO (0.53) (Fig. 5e). All methods with output except PENN and DCPR were above the 95th percentile of their permutation null (0.21–0.24) for amplitude reconstruction. The cohort has three recorded covariates: sequencing run, sex, and breeding generation. We adjusted for them in three ways: ComBat-seq correction of the counts for sequencing run, with sex and generation retained as model covariates (all methods), regression of all three out of the expression matrix (CHIRAL), and CYCLOPS 2.0’s native covariate mode, which received all three as categorical covariates. Against the separate liver reference, ComBat-seq raised the median acrophase error for eight of nine methods (by 0.03–0.82 h) and lowered it only for Cyclum (-0.58 h), whereas CYCLOPS 2.0’s batch mode lowered it by 1.69 h and CHIRAL’s covariate regression by 0.41 h, each relative to the unadjusted run (Fig. 5f). ComBat-seq changed the amplitude correlation by 0.10 for COFE and -0.15 for TEMPO, and CYCLOPS 2.0’s batch mode raised it by 0.29 (Fig. 5g).

## Discussion

Performance varied widely between methods run under a common protocol. CHIRAL had the lowest mean aligned MAE, our primary outcome (1.80 h), followed by TEMPO, CYCLOPS and CYCLOPS 2.0 (2.49-2.80 h). The other six methods had mean errors of 3.91-4.86 h, against a chance level of 5.08-5.23 h. The ordering of the leading methods remains uncertain, because the benchmark contains only six study families and the main runs use one random seed: post-hoc comparisons separated CHIRAL from COFE, PENN and PROTECT, but not from TEMPO, CYCLOPS, or CYCLOPS 2.0.

A principal-component baseline built from twelve known clock genes had a mean error of 2.15 h, lower than every inference method except CHIRAL, and a baseline using just two genes was similar at 2.20 h. The baselines show how much accuracy prior knowledge of core clock genes provides in datasets where those genes cycle, and they also depend on that cycling: across the 25 datasets their errors were associated with clock-gene rhythm strength (Spearman ρ = -0.94 for clock-gene PCA and -0.67 for two genes). The leading methods (CHIRAL, TEMPO, CYCLOPS, CYCLOPS 2.0) all used circadian prior information, but this was not sufficient for high accuracy since DCPR also used a circadian seed-gene list and had a higher mean error. The four methods without circadian prior information (COFE, Cyclum, PENN, PROTECT) also had higher mean errors than the leading four.

Not all of the methods were developed for the data used here. reCAT and Cyclum were developed to order single cells along the cell cycle, PROTECT for proteomic data and TEMPO for circadian phase in single-cell counts, whereas this benchmark applies every method to bulk RNA-seq samples collected around the day. The rank of a method here therefore reflects how well it transfers to this setting, together with its model and the configuration we ran.

Accuracy was associated with clock-gene rhythm strength (median error over methods, Spearman ρ = -0.58). On the human striatal datasets no method had an error below 2.9 h, and mean errors were lower in simulations than in real data for every method. The simulations also changed the order of the methods: CYCLOPS, COFE and CYCLOPS 2.0 reached errors of 0.3-0.4 h in the reference simulation, while their errors on real data were larger. The simulated model may favor the assumptions of some methods, so accuracies in simulations should not be read as accuracies on real data. They are, however, useful for isolating single factors such as the sampling design, or number of samples that may affect a tool’s performance. In the positive-control simulation, designed so that the clock is easy to find, eight of ten tools had lower errors than both baselines (0.19-0.98 h), while PROTECT and PENN did not (2.08 and 2.27 h). We also could not reproduce their published results either (Supplementary Table S1), so these findings pertain to the performance of these tools run with our configuration.

The random seed matters for some methods. For methods with substantial seed-dependent variation, small differences from a single run may not persist across repeated runs. The seed-to-seed standard deviation of the aligned MAE was 0.42–0.52 h for five methods (DCPR, PROTECT, TEMPO, PENN and Cyclum), comparable to differences between adjacent methods in the top group, whereas it was 0.01 h for CYCLOPS and 0.07 h for CHIRAL. CYCLOPS 2.0 gave identical output at every seed setting because its fitting routine resets the random seed internally, so its lack of variation is not a measure of stability. Reporting performance across several initializations is therefore important for methods that depend on their initialization.

On the ten datasets with batch labels, ComBat-seq correction lowered the mean error of eight of the ten methods, most for CYCLOPS (-1.98 h), COFE (-1.08 h), reCAT (-0.90 h) and CYCLOPS 2.0 (-0.89 h). It left CHIRAL unchanged (1.83 and 1.84 h) and raised the error of TEMPO (+0.28 h) (Fig. 2d, Supplementary Fig. S12). Adjustment can, however, remove circadian signals when a covariate is associated with time. In the four GSE202537 post-mortem datasets, the binned post-mortem interval alone explained 0.61-0.64 of the combined variation in the cosine and sine of the reference time (unadjusted R squared with three parameters, Supplementary Fig. S8). We regressed the covariates out of each gene in CHIRAL’s input matrix, the adjustment its authors applied in their GTEx analysis, and ran CHIRAL and the two clock-gene baselines on the adjusted matrix. On these four datasets the adjustment raised CHIRAL’s mean error from 3.56 to 5.44 h and the baselines’ from 3.96 to 5.39 h (clock-gene PCA) and from 4.35 to 5.18 h (two genes), beyond the chance level of about 5 h, and no dataset improved. Because the baselines use only the clock genes and lost accuracy in the same way, this suggests that the adjustment removed time-associated expression from those genes, rather than affecting CHIRAL alone.

Diversity Outbred datasets test whether the ranking from the labelled datasets carries over to a large cohort without collection times, and for the leading method it did: CHIRAL was the only method consistent with both liver references. For adipose, however, no method agreed with the reference. However, this could either be the methods’ failing, or the fact that we are using a reference from a different study, and we cannot tell which. Agreement of the inferred phases between the tissues, however, does not depend on an external reference, since liver and adipose from the same mouse should give the same time. Yet, even CHIRAL reached a correlation of only 0.47, so substantial error remains in the best case. For four methods the agreement is also not independent of the reference: the seed lists of CYCLOPS, CYCLOPS 2.0 and DCPR and the acrophase priors of TEMPO are derived from the Zhang atlas, the same atlas used as reference (open circles in Fig. 5a,b). For these four methods agreement with the Zhang atlas is partly built in, and only the second liver reference, which excludes the Zhang studies, tests them independently.

On these results we recommend CHIRAL as the first choice for bulk RNA-seq data: it had the lowest mean error and ranked among the four best methods in every dataset. However, it could not be statistically separated from TEMPO, CYCLOPS and CYCLOPS 2.0, and its median advantage over them within a dataset was 0.2 - 1.1 h. Hence these three are reasonable alternatives.

The study has limitations. The comparison rests on 25 datasets from 6 independent study families, and each simulated condition was generated once. Reference times for the GSE202537 post-mortem datasets are uncertain, since unwitnessed deaths were assigned a time-of-death window of generally 2 h and occasionally 4 h. We ran each tool at one predefined configuration taken from its publication or documentation where these specify it and from the software defaults otherwise, and some tools were designed for other data (single-cell, cell-cycle, proteomic), so the results are not evaluations of those tools for their intended use.

## Methods

### Method selection

Methods were selected that infer the circadian phase of every sample in an unlabelled bulk expression matrix. Supervised methods that train on a labelled dataset were not included. We identified 10 methods for evaluation: CYCLOPS (Anafi et al. 2017), CYCLOPS 2.0 (Li et al. 2024), CHIRAL (Talamanca et al. 2023), TEMPO (Auerbach et al. 2022), COFE (Ananthasubramaniam and Venkataramanan 2025), DCPR (Han et al. 2026), PENN (Ansary Ogholbake and Cheng 2023, 2024), PROTECT (Ansary Ogholbake and Cheng 2025), Cyclum (Liang et al. 2020), and reCAT (Liu et al. 2017). See Methods Table 3 for details. One method, ORI (Larriba et al. 2020) was not evaluated since there was no public documentation or example code and it failed on our attempts to run it. Another method, ESOCVD, has a preprint (Ji et al. 2020) but no publication and was not evaluated.

### Dataset selection

We selected a total of 20 published datasets with true reference times in either mouse or human tissues, see Methods Table 1. We further selected 2 mouse datasets without reference times. See Supplemental Methods for details.

### Simulation datasets

We simulated 13 datasets using dependentsimr (Yang et al. 2025), referencing real datasets. In short, negative-binomial marginals, a mean *q*_*g*_ and dispersion α_*g*_ per gene, were estimated with the DESeq2 estimator (Love et al. 2014) on GTEx bulk liver counts (GTEx Consortium 2020), and dependence between genes was represented by a Gaussian copula whose covariance was estimated with dependentsimr’s spiked-Wishart method. For gene *g* and sample *i* at time *t* _*i*_ the mean expression was set to be

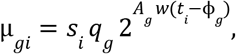

where *w* is a periodic waveform (a cosine except in the waveform tests) and ϕ_*g*_ the acrophase. *A*_*g*_ is the amplitude on the log2 scale. Which genes are rhythmic, and their ϕ_*g*_ and *A*_*g*_, were taken from a MetaCycle (Wu et al. 2016) rhythmicity call (p < 0.05) on the GSE151565 mouse liver or hypothalamus time course orthologous. Size factors *s*_*i*_ were resampled from GTEx library sizes scaled to mean 1. Batch effects, where present, multiply µ_*gi*_ by an exponential factor of the form 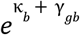, where b denotes the batch of sample I (here, κ _*b*_ denotes the global shift in expression across the batch, whereas 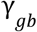 is the per-gene shift). The parameters for the distributions of κ _*b*_ and 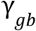 were determined empirically from mouse studies.

Simulations used 25,972 (liver) and 26,193 (hypothalamus) genes, of which 7,047 and 2,659 are rhythmic, respectively. Simulations had as baseline 100 samples over one cycle every 2 h, dispersion scaled to 0.15, no batch effect, random seed 17. We also simulated variations that change one setting at a time, see Methods Table 2 and Supplemental Methods. Both tissues use GTEx liver marginals, so the hypothalamus simulation combines hypothalamic rhythms with liver expression levels.

### Preprocessing

Preprocessing steps were selected for each method according to their published paper, or their documentation and example code if not specified in the publication. Datasets were obtained as raw counts and were converted, for each method, as TPM or FPKM values, and optionally log transformed, z-scored and/or low counts removed. Methods were provided with any additional methods (lists of known rhythmic genes, acrophase priors) as recommended by their authors. See Supplemental Methods for details.

### Evaluation criteria

Aligned circular mean absolute error (MAE) was the primary performance measure and was used for method rankings. Circular correlations and gene-level rhythm reconstruction measures were reported as secondary outcomes. In datasets without known rhythms gene-level rhythm recovery was the primary outcome.

### Aligned circular mean absolute error

For inferred phases 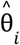 and reference phases θ_*i*_, *i* = 1, …, *n*, let *d*(*a, b*) = *arccos*(*cos*(*a* − *b*)) be the shortest angular distance. The aligned circular mean absolute error is defined as

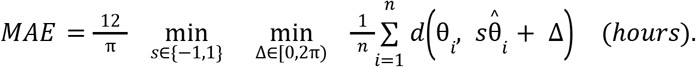

MAE is in the units of hours, ranges from 0 to 6hr, and lower values represent better reconstruction of phases.

### Circular correlation

We use absolute Fisher–Lee circular correlation (Fisher and Lee 1983):

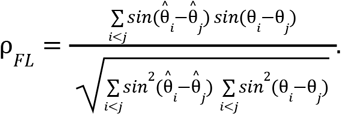

Since inferred time orientation is arbitrary, we report the absolute value, which ranges from 0 to 1 with higher values representing better reconstruction of phases.

### Circular rank correlation

A circular rank correlation was also computed, as the same Fisher-Lee correlation applied to circular ranks 2π*r*_*i*_ /*n* of each set of phases (phases ranked as *r*_*i*_, ties given their average rank); its absolute value is reported.

### Reconstructed rhythms

Methods were evaluated on the ability for the inferred phases to reconstruct rhythms of genes. In datasets with reference times available, reference rhythms were computed by cosinor fits of *log*_2_(*TPM* + 1) expression by reference time. Reconstructed rhythms were defined as cosinor fits to the same expression values but using the inferred sample phases instead. Genes were considered reference-rhythmic if they had Benjamini-Hochberg q-value less than 0.05 and had an acrophase standard error of at most 1h. Gene acrophase refers to the peak time of a gene; amplitude to the *log*_2_-scale cosinor amplitude (i.e., half the peak-to-trough). These were summarized as: mean MAE between reconstructed gene acrophases and reference acrophases; Spearman correlation of reconstructed amplitude to reference amplitude; and the ROC AUC of binary rhythmicity.

### Ranking

Methods were ranked by the mean aligned circular MAE with equal dataset weights in the main accuracy comparison. If any compared method had a missing or undefined value, that dataset was excluded for all methods in that test.

## Data and code availability

Real expression datasets were obtained from the GEO accessions listed in Methods Table 1. Benchmark code, simulation code, and configurations are available at https://github.com/tgbrooks/circ_infer_benchmark.

## Acknowledgements

SP received funding support from the National Heart Lung and Blood Institute (T32HL170968). GRG and TGB received funding support from the National Center for Advancing Translational Sciences Grant (UL1TR001878).

## Disclosures

No authors of any methods had any input on this analysis. Claude Opus 5.5 was used in generating the code, figures, and drafting this manuscript.

## Notes

### Competing Interest Statement

The authors have declared no competing interest.

