## Supplementary Information for "A Benchmark of Unsupervised Circadian Phase Inference Methods for Bulk Transcriptomic Data"

#### 1. Supplementary Methods

##### 1.1 Study design

Overview. We evaluated 10 methods that infer the circadian phase of every sample in an unlabelled bulk expression matrix. Each method was run using a single predefined configuration for each dataset, specifying the preprocessing procedure, parameter settings and, for two methods, a fixed fallback rule. These settings were based on the authors' publications and code, supplemented by their recommendations where available, and constrained by each method's input requirements and model assumptions. All settings were fixed before evaluation. Methods do not report phases in a common convention: the origin of the cycle and its direction differ between them, and only some anchor these with clock-gene priors. Inferred phases were therefore compared with recorded collection times after fitting a rotation and a reflection. Because those two parameters are fitted using the reference times, the evaluation assesses how well each method reconstructs the relative phase positions of samples, including the spacing between them, without evaluating the absolute origin or direction of the cycle. Method rankings were based on the predefined configurations described above.

Sensitivity analyses. We assessed the effects of input normalization, random seed, seed gene list, tissue-matched seeding, and batch and covariate adjustment by varying one factor at a time. These analyses used the same underlying datasets and did not contribute to the method rankings. Two modes of a ranked method that differ from it in a single input, CYCLOPS 2.0 given batch labels and PENN given a seed list, are compared with their base method only.

Excluded methods. An eleventh method, ORI, was wrapped but is not evaluated. It has no public repository, no example data and no published accuracy figure, so the wrapper could not be checked against any reference, and it failed on most runs. Its coverage is reported as a limitation only. Another method, ESOCVD has a public repository and preprint but has not been published and was not evaluated.

##### 1.2 Datasets

Collection. We assembled 36 datasets, of which 35 were run through every method: 22 real and 13 simulated. A further mouse liver atlas (1,096 samples from 57 studies) was used only to calibrate the simulations and as a separate reference map. Two of the run datasets, liver and adipose from a Diversity Outbred mouse cohort (Tyler et al. 2025), have no recorded collection times, leaving 33 with reference times. Of these, 25 datasets (16 real and 9 simulated) were selected for the main accuracy comparison. The remaining 8 datasets were used as follows:

1. 4 challenge datasets with an absent or weak circadian clock: two overlapping nucleus accumbens sets from GSE202537 (Ketchesin et al. 2023) with little detectable clock-gene rhythm, and Bmal1$^{-/-}$ and Per1/Per2 double-knockout livers sampled in constant darkness (Aviram et al. 2021);

2. one positive control, a simulation built so that the rhythm is easy to reconstruct;
3. 3 waveform tests, in which the simulated cosine is replaced by another shape.

Methods Table 1. Real datasets with recorded collection times. Sampling is computed from each dataset's timepoints.

| Dataset | Organism | Tissue | N | Sampling | Time label | Role |
|---|---|---|---|---|---|---|
| GSE151565 | mouse | liver | 77 | 3 h grid, 8 phases (raw span 0–36 h) | controlled protocol | ranked |
| GSE171975 | mouse | liver | 48 | 4 h grid, 6 phases (raw span 0–44 h) | constant darkness (CT) | ranked |
| GSE108539 | human | muscle | 57 | 4 h grid, 6 phases | controlled protocol | ranked |
| GSE202537_Caudate | human | brain (caudate) | 36 | continuous, one time per subject | postmortem TOD, ±1 h | ranked |
| GSE202537_Putamen | human | brain (putamen) | 36 | continuous, one time per subject | postmortem TOD, ±1 h | ranked |
| GSE202537_Caudate_all | human | brain (caudate) | 72 | continuous, one time per subject | postmortem TOD, ±1 h | ranked |
| GSE202537_Putamen_all | human | brain (putamen) | 72 | continuous, one time per subject | postmortem TOD, ±1 h | ranked |
| GSE201207_adrenal | mouse | adrenal | 36 | 4 h grid, 6 phases | controlled protocol | ranked |
| GSE201207_heart | mouse | heart | 36 | 4 h grid, 6 phases | controlled protocol | ranked |
| GSE201207_hypothalamus | mouse | brain (hypothalamus) | 36 | 4 h grid, 6 phases | controlled protocol | ranked |
| GSE201207_kidney | mouse | kidney | 36 | 4 h grid, 6 phases | controlled protocol | ranked |
| GSE201207_lung | mouse | lung | 36 | 4 h grid, 6 phases | controlled protocol | ranked |
| GSE201207_muscle | mouse | muscle | 36 | 4 h grid, 6 phases | controlled protocol | ranked |
| GSE151565_Cortex | mouse | brain (cortex) | 78 | 3 h grid, 8 phases | controlled protocol | ranked |
| GSE151565_Hypothalamus | mouse | brain (hypothalamus) | 77 | 3 h grid, 8 phases | controlled protocol | ranked |
| GSE151565_Striatum | mouse | brain (striatum) | 60 | 3 h grid, 8 phases | controlled protocol | ranked |
| GSE202537_NAc | human | brain (accumbens) | 36 | continuous, one time per subject | postmortem TOD, ±1 h | challenge (weak/absent clock) |
| GSE202537_NAc_all | human | brain (accumbens) | 71 | continuous, one time per subject | postmortem TOD, ±1 h | challenge (weak/absent clock) |
| GSE171975_Bmal1KO | mouse | liver | 46 | 4 h grid, 6 phases (raw span 0–44 h) | constant darkness (CT) | challenge (weak/absent clock) |
| GSE171975_PerDKO | mouse | liver | 46 | 4 h grid, 6 phases (raw span 0–44 h) | constant darkness (CT) | challenge (weak/absent clock) |

Real data. Count matrices and sample annotations were obtained from the Gene Expression Omnibus: mouse liver, cortex, hypothalamus and striatum (GSE151565), mouse liver from wild-type and clock-mutant animals (GSE171975) (Aviram et al. 2021), human skeletal muscle (GSE108539) (Perrin et al. 2018), human caudate, putamen and nucleus accumbens (GSE202537) (Ketchesin et al. 2023), and six tissues from mice of three ages (GSE201207) (Wolff et al. 2023). Each tissue is a separate dataset. GSE202537 is used twice per region, as control subjects only and as the full cohort including subjects with psychosis. Recorded collection times were converted to phase as $\theta = 2\pi\,(t\ mod\ 24)/24$. These times record when a sample was taken, not each individual's internal circadian phase.

Dependence between datasets. The 16 real datasets in the main accuracy comparison come from 5 GEO series. Tissues of one series were taken from the same animals or donors, and the control-only and full-cohort versions of GSE202537 share subjects. All simulations share one generator, calibration and random seed. We therefore treat each GEO series, and the simulations as a whole, as a study family, and use families rather than datasets as the unit of resampling and testing (Section 1.5.3). The families are not fully independent either: the simulations take their rhythm template from GSE151565. Every family-level test is therefore also reported for the real-data families alone.

Repeated subjects. GSE108539 contains serial muscle biopsies from 10 volunteers, so its samples are not independent donors. Donor identity was taken from the GEO sample titles and used only during evaluation: in permutation tests (Section 1.5.2) and in every gene-level cosinor fit (below, Section 1.5.5). No method received it. Every other dataset has one sample per animal or donor.

Reference times. Recorded sampling times were used as reference times for experimental datasets, and recorded times of death were used for human post-mortem datasets. For GSE202537 time-of-death values are up to 4 h off.

Independence. GSE108539 was used in the development of CHIRAL (Talamanca et al. 2023); it is kept in the main accuracy comparison and its effect is tested by removal.

Clock-gene rhythm $R^2$. Each dataset was characterised by how strongly twelve canonical clock genes (ARNTL, CIART, CRY1, CRY2, DBP, NPAS2, NR1D1, NR1D2, PER1, PER2, PER3, TEF) cycle against the reference times. For each gene we fitted $y = \beta_0 + \beta_1 cos\omega t + \beta_2 sin\omega t$, $\omega = 2\pi/24$, to $log_2(TPM + 1)$, with a fixed intercept per donor for GSE108539, and recorded the partial $R^2$ of the rhythm terms and the amplitude $\sqrt{\beta_1^2 + \beta_2^2}$ in log2 units. The dataset value is the median over genes.

##### 1.2.1 Simulated datasets

Methods Table 2. Simulated datasets. Every simulation except the positive control differs from the reference configuration in one setting; the positive control changes several (see text).

| Dataset | Tissue | N | Sampling | Role |
|---|---|---|---|---|
| liver-none-N100-morton | liver | 100 | 2 h grid, 12 phases | ranked |
| hypothalamus-none-N100-morton | hypothalamus | 100 | 2 h grid, 12 phases | ranked |
| liver-balanced-N100-morton | liver | 100 | 2 h grid, 12 phases | ranked |
| liver-mild-N100-morton | liver | 100 | 2 h grid, 12 phases | ranked |
| liver-none-daytime-N100-morton | liver | 100 | continuous, 95/100 within a 12 h window | ranked |
| liver-none-window-N100-morton | liver | 100 | continuous, 11.9 h span | ranked |
| liver-none-clustered-N100-morton | liver | 100 | continuous, one time per subject | ranked |
| liver-none-N24-morton | liver | 24 | 2 h grid, 12 phases | ranked |
| liver-none-N48-morton | liver | 48 | 2 h grid, 12 phases | ranked |
| liver-none-N100-morton-clean | liver | 100 | 2 h grid, 12 phases | positive control |
| liver-none-N100-morton-square | liver | 100 | 2 h grid, 12 phases | waveform stress |

| Dataset | Tissue | N | Sampling | Role |
|---|---|---|---|---|
| liver-none-N100-morton-sawtooth | liver | 100 | 2 h grid, 12 phases | waveform stress |
| liver-none-N100-morton-spline | liver | 100 | 2 h grid, 12 phases | waveform stress |

Generative model. Counts were simulated with dependentsimr (Yang et al. 2025). Negative-binomial marginals, a mean $q_g$ and dispersion $\alpha_g$ per gene, were estimated with the DESeq2 estimator (Love et al. 2014) on GTEx bulk liver counts (GTEx Consortium 2020), and dependence between genes was represented by a Gaussian copula whose covariance was estimated with the package's spiked-Wishart method. The rhythm was applied to the negative-binomial mean before counts were drawn. For gene $g$ and sample $i$ at time $t_i$ the mean was

$$\mu_{gi} = s_i q_g 2^{A_g w(t_i - \phi_g)},$$

where $w$ is a unit periodic waveform with range $[-1, 1]$ (a cosine except in the waveform tests) and $\phi_g$ the acrophase. $A_g$ is the amplitude on the log2 scale: it was taken directly from MetaCycle's amplitude estimate, which was computed on $log_2(TPM + 1)$. Writing $a_g = 2^{A_g} - 1$, the mean at peak is $(1 + a_g) s_i q_g$ and at trough $s_i q_g/(1 + a_g)$, so $a_g$ is the elevation of the peak over $q_g$, not a symmetric linear-scale amplitude over a mesor. Size factors $s_i$ were resampled from GTEx library sizes scaled to mean 1, dispersions were multiplied by a constant scale, and each sample was drawn from the copula with its own means. Batch effects, where present, multiply $\mu_{gi}$ in the same way. Which genes are rhythmic, and their $\phi_g$ and $A_g$, were taken from a MetaCycle (Wu et al. 2016) rhythmicity call ($p < 0.05$) on the GSE151565 liver or hypothalamus time course; mouse symbols were matched to human symbols by upper-casing, which is a match of names that assumes, without checking, that the genes are orthologous.

Dispersion. Fitting the same estimator to the real datasets of this benchmark, with the rhythm in the model so that it is not counted as noise, gave median dispersions 0.084–0.222 times that of GTEx liver; 0.15 lies near the middle of that range.

Time sampling and sample size. Four simulations change the sampling times at N = 100: uniform spreads the 100 samples evenly over the twelve time points of a 2h grid (8 or 9 samples per time point), daytime places 94 of 100 samples between 08:00 and 18:00 (Methods Table 2 reports the densest 12 h window), window samples continuously over 12 h from 08:00, and clustered places samples in three groups of 33 or 34 centered on 00:00, 08:00, and 16:00. Two reduce N to 48 and 24 on the same 2 h grid.

Batch effects. Two simulations assign samples to four batches, either independently of time (balanced) or partly confounded with it (mild). Batch magnitude was estimated from 17 control studies (304 samples, 10,554 genes) of the mouse liver atlas, separating within- from between-laboratory variation; the within-laboratory estimate was applied as a per-gene log2 fold-change SD of 0.476 on 50.7% of genes (Laplace-distributed) and a per-batch library-depth SD of 0.109 (natural log). This estimate rests on five degrees of freedom from four laboratories, in liver only.

Positive control. The positive control changes four settings of the reference: every $A_g$ doubled, the rhythmic fraction raised to 0.3, twelve core clock genes given $A_g = 3$ (an 8-fold peak over $q_g$) at acrophases measured in mouse liver, and dispersion lowered to 0.1.

Waveform tests. Three simulations replace the cosine with a square wave (duty cycle 0.5), a sawtooth rising for 4.8 h and falling for 19.2 h, or per-gene shapes from spline fits from the mouse liver atlas, each peak-aligned and scaled to the range [−1, 1]. The first harmonic carries 81% of the variance of the square wave and 83% of the sawtooth. Matching the range does not match signal variance (over a cycle the unit-range square wave has twice, and the sawtooth two-thirds, the variance of the cosine), so these tests change signal power as well as shape.

#### 1.3 Methods evaluated

Methods Table 3. Methods evaluated, with the data type each was designed for and its authors' preprocessing and accuracy measure, as stated in each method's paper and documentation.

| Tool | Designed for | Authors' preprocessing | Authors' accuracy measure |
|---|---|---|---|
| CYCLOPS (Anafi et al. 2017) | bulk transcriptome | top 10k probes, extremes truncated, $S_{ij} = (X_{ij} - M_i)/M_i$, eigengenes | median absolute error, circular correlation |
| CYCLOPS 2.0 (Li et al. 2024) | bulk, with covariates | as CYCLOPS, plus covariate terms in the model | error on synthetic data |
| CHIRAL (Talamanca et al. 2023) | bulk transcriptome | genes with mean count < 10 removed, TMM, $log_2(CPM + 0.25)$, per-gene standardization | circular correlation, mean / median absolute error, RMSE |
| COFE (Ananthasubramaniam and Venkataramanan 2025) | bulk and multi-omic | $log_2$ CPM, features with mean $\geq 7$, standardization | median absolute position error |
| DCPR (Han et al. 2026) | bulk transcriptome | z-normalization by the user | AUC of the error CDF, median absolute error, circular correlation |
| PENN (Ansary Ogholbake and Cheng 2023, 2024) | bulk transcriptome | per-gene z-score | normalized AUC |
| TEMPO (Auerbach et al. 2022) | single-cell, circadian | raw UMI counts | posterior calibration, likelihood against shuffled phases |
| PROTECT (Ansary Ogholbake and Cheng 2025) | proteomics | per-protein z-score | normalized AUC |
| Cyclum (Liang et al. 2020) | single-cell cell cycle | $log_2(x + 1)$, then z-score | cell-cycle classification accuracy |
| reCAT (Liu et al. 2017) | single-cell cell cycle | genes with mean TPM < 2 removed, $log_2(TPM + 1)$ | agreement with known cell-cycle stage |

Scope. reCAT and Cyclum were developed to infer cell-cycle progression from single-cell expression data, with reCAT using a predefined list of cell-cycle genes. PROTECT was developed for proteomic data. We evaluated these three methods for circadian phase inference from bulk RNA-seq data. TEMPO was developed for circadian phase inference from single-cell UMI counts. Its count likelihood accepts bulk RNA-seq counts, although these data differ in depth and dispersion from those for which the method was developed.

PENN. The PENN repository implements the 2023 conference method (Ansary Ogholbake and Cheng 2023) (z-scoring, four principal-component eigengenes and an autoencoder with a cosine-fitting loss), not the 2024 journal method (Ansary Ogholbake and Cheng 2024), whose code was not released. All PENN results refer to the 2023 method.

##### 1.3.2 Baselines

Two simple deterministic baselines were scored on the same terms as the methods, on linear TPM, with the matrix and gene order fixed before scoring:

- Clock-gene PCA: the twelve clock genes were z-scored and each sample's phase taken as $atan2(PC2, PC1)$;
- Two genes: $atan2$ of two z-scored clock genes, using the first available pair of ARNTL/NR1D1, NPAS2/DBP, CRY1/DBP, PER2/NR1D1.

###### 1.3.3 Execution and coverage

Each method ran in its own isolated software environment, built from the authors' environment where one is published, with the method's code unchanged. A failed run was retried at most twice with identical settings.

##### 1.4 Preprocessing and method configuration

Gene identifiers. Ensembl gene or transcript identifiers were mapped to gene symbols with Ensembl annotation tables for each organism (BioMart, downloaded April 2026; the release was not pinned). Identifiers without a symbol were dropped. Where several rows mapped to one symbol, the row with the largest total count was kept. GSE171975 is deposited as gene-symbol rows plus 3,572 Ensembl transcript rows; none of those transcript identifiers is present in the current annotation, so they were dropped (4.7% of counts). Clock-gene aliases (BMAL1 → ARNTL, CHRONO → CIART, REVERBA → NR1D1 and others) were mapped to their corresponding gene symbols. Gene symbols were matched case-insensitively, allowing mouse gene lists to be used with human datasets where symbols matched. Rows lost at each step were logged per dataset.

Normalization. We computed TPM and FPKM from counts and gene lengths (the longest annotated transcript per symbol, an approximation to the effective length of a gene-level count; genes without a length are absent from these two matrices) and TMM-normalized counts per million with edgeR (Robinson and Oshlack 2010). Each was also log-transformed as $log_2(x + c)$, with $c$ taking each of the values 10, 1, 0.25, 0.1. The 0.25 offset is CHIRAL's pseudocount. With raw counts this gives 16 matrices per dataset.

Method-specific filters. Three methods' authors remove lowly expressed genes, and these filters were applied as separate input matrices:

- CHIRAL: genes with mean count below 10 removed before TMM normalization; then $log_2(x + 0.25)$
- COFE: genes with mean $log_2(CPM + 1)$ below 7 removed; then $log_2(x + 1)$;
- reCAT: genes with mean TPM below 2 removed; then $log_2(x + 1)$.

This gives 19 input matrices per dataset. Matrices were stored rounded to three decimal places, which is the precision every method received.

###### 1.4.1 Method preprocessing configurations

All method configurations were fixed before evaluation. Input selection followed each method's model requirements: TEMPO received integer counts for its count likelihood, while CHIRAL and reCAT received log-transformed expressions because they require this input but do not perform the transformation internally. Otherwise, we followed the authors' documented procedures or code defaults. TPM was used when neither the model requirements nor the documentation specified an input format. No configuration was chosen by comparing scores on the benchmark datasets.

We followed the authors' procedures for CHIRAL's gene filtering, normalization, pseudocount and standardization; COFE's and reCAT's gene filtering; and DCPR's z-normalization. For PENN, we used the per-gene z-scoring specified in the publication; its code instead applies a single matrix-wide scaling.

Methods Table 4. Input matrix each method receives on every dataset.

| Tool | Input and preprocessing steps | Extra data provided |
|---|---|---|
| CHIRAL | log2(TMM-CPM + 0.25), TMM computed after removing genes with mean count < 10 | 12 core clock genes |
| COFE | log2(TMM-CPM + 1), genes with mean log2 CPM < 7 removed | |
| CYCLOPS | TPM | 31 genes rhythmic in 75% of mouse tissues or tissue-specific cyclers if available |
| CYCLOPS 2.0 | TPM | 34 ubiquity rhythmic genes unioned with tissue-specific cyclers |
| Cyclum | log2(TPM + 1), z-scored | |
| DCPR | TPM, z-scored | Tissue-specific rhythmic genes if available |
| PENN | TPM, z-scored | |
| PROTECT | TPM, z-scored | |
| reCAT | log2(TPM + 1), genes with mean TPM < 2 removed | |
| TEMPO | raw counts | 26-gene clock list, acrophase prior |

Parameters. Methods ran at their defaults except for the following, taken from the authors' published settings: CYCLOPS with seed-gene CV bounds 0.14–0.70, blunting 0.975 and the ten best of its restarts; CYCLOPS 2.0 with at most six eigengenes, seed-gene CV at most 0.7, eigengene contribution threshold 0.03 and no training on collection times; PROTECT with layer widths scaled to the number of features as its paper describes; and reCAT on one core, because its parallel path is not reproducible under a fixed seed.

Random seeds. Every stochastic method used one seed for every run: 12345 for CYCLOPS and CYCLOPS 2.0 (their scripts' default) and 42 for all others. A run that failed at that seed was not rerun at another.

Extra data provided. Five methods take additional data, such as a list of core clock gene identifiers or known rhythmic genes. We used each method's published datasets for these, see Methods Table 4 for details.Tissue-specific rhythmic genes were taken from  the Zhang mouse atlas (Zhang et al. 2014) when available.

Fallback rules. For two methods the predefined configuration is a sequence of attempts, triggered only by an event observable without reference times:

- TEMPO abstains when its fit is no better than shuffled phases. It was given an ordered list of acrophase priors and moved to the next only on abstention: for the 17 liver datasets the liver clock prior, then the core clock and ubiquitous-cycler prior, then none; for all others the core prior, then none, then the liver prior. Any other error ended the sequence.
- DCPR without a usable tissue list runs in its authors' no-list mode, in which all genes are passed and DCPR keeps the 1,000 most rhythmic by its own cosinor fit. This applies to 11 datasets, three of them because DCPR failed on the tissue list.

The prior or mode used and the number of attempts are recorded for every run, and TEMPO's abstention is reported separately from execution failure.

Run outcomes. Each scheduled run of a method on a dataset had one of four outcomes:

1. scored: a phase for each sample;
2. abstained: the method itself declined to return phases (TEMPO only);
3. failed: no output after retries;
4. collapsed: output in which samples are concentrated on a very small number of phases (flagged when there are at most 2 distinct phases).

All runs that returned finite phase estimates for the required samples were included in accuracy evaluation, including runs with collapsed outputs. Collapsed outputs were flagged and scored using the same aligned circular MAE as other outputs. Runs that failed, abstained or returned unusable output were excluded from accuracy calculations and reported separately. A secondary measure that is undefined for a run does not remove that run's MAE.

#### 1.5 Evaluation

##### 1.5.2 Permutation tests

For each run, inferred phases were permuted and the MAE relative to the (unpermuted) reference phases were recomputed. Let $MAE_{obs}$ denote the observed aligned error and $MAE_b$ the aligned error from permutation $b$. With $B$ = 2,000 permutations, the one-sided permutation p value was calculated as

$$p = \frac{1 + \sum_{b=1}^{B} 1\{MAE_b \le MAE_{obs}\}}{B+1}.$$

Collection-time concordance was considered statistically significant when the one-sided permutation p value was below 0.05. Because the permutations keep the method's phase distribution and the dataset's sampling times, the mean shuffled error depends on both, as well as on $n$; it is therefore estimated per run rather than set to a constant. The same permutations give the permutation mean and p value for $|\rho_{FL}|$. Per method we report the fraction of datasets with significant concordance.

For GSE108539, phases were permuted only among samples of the same donor. Within-donor permutation assumes that a donor's samples are exchangeable with respect to time under the null hypothesis. It accounts for donor-level differences but not for serial correlation between a donor's successive biopsies.

##### 1.5.3 Ranking

Point summaries. For each method and metric, mean and median performance were reported across eligible datasets using the available dataset-level values. Unless otherwise specified, each dataset received equal weight regardless of sample size. Family-weighted means and medians were also calculated, assigning equal total weight to each study family and dividing that weight equally among its datasets with available values for the corresponding method and metric. Summaries were reported across all eligible datasets and separately for real and simulated datasets.

Methods were ranked by the mean aligned circular MAE with equal dataset weights in the main accuracy comparison. For summaries comprising multiple study families, 95% confidence intervals were estimated by resampling whole study families with replacement, retaining all datasets within each sampled family and recalculating the corresponding summary.

Rank tests. For the Friedman test and Nemenyi post-hoc comparisons (Demšar 2006), related datasets were grouped into study families, with real datasets grouped by GEO study and simulated datasets treated as a separate family. These tests were applied to aligned circular MAE only; secondary measures were summarized descriptively. A dataset was included only if values were available for every method being compared. If any compared method had a missing or undefined value, that dataset was excluded for all methods in that test, while its available results were retained in the reported performance summaries. Within each family, method means were calculated from the remaining common datasets. A family was retained if at least one such dataset remained; otherwise, it was excluded from the test. Methods were ranked within each retained family, and these ranks were compared using the Friedman test and Nemenyi post-hoc comparisons. This restriction applied only to these tests; reported performance summaries and descriptive rankings used each method's available values. The tests were run for all families and for the real-data families alone. With this few blocks the usual chi-square approximation is unreliable (Demšar 2006), so significance was taken from 10,000 permutations in which method labels are shuffled independently within each family. Nemenyi comparisons rely on a large-sample approximation and are reported as descriptive.

Rank stability. Rank stability was assessed using 5,000 bootstrap resamples of whole study families. Each resample was formed by selecting the original number of study families with replacement and retaining all datasets within each selected family. The mean aligned circular MAE of each method was recalculated using its available dataset-level values with equal dataset weights, and methods were ranked from lowest to highest error. The frequency with which each method ranked first was reported. Given the small number of study families, these frequencies were interpreted as measures of ranking stability within the benchmark. Rankings were also recalculated after omitting each study family in turn and within groups defined by tissue, organism and the source of reference times. The analysis excluding GSE108539 assessed sensitivity to inclusion of a dataset used to benchmark CHIRAL and compare seed gene lists in its original publication (Talamanca et al. 2023).

##### 1.5.4 Positive control and challenge datasets

Performance on the positive-control dataset was assessed using aligned circular MAE. A method met the positive-control criterion if its MAE was below 1.5 h. This threshold was fixed before the inference methods were evaluated, above the larger of the two baseline errors measured when the dataset passed its quality-control checks (0.73 h and 1.10 h). Those checks applied the baselines to $log_2(count + 1)$ of the clock genes; the benchmark applies them to linear TPM (Section 1.4.2), where they score 1.01 h and 1.60 h. The prespecified threshold was kept. Circular correlations and gene-level rhythm reconstruction measures were also reported as secondary outcomes.

The challenge datasets were used to assess phase inference when clock-gene rhythms were weak or disrupted. Weak clock-gene rhythms do not establish an absence of time-dependent expression; for example, rhythms of approximately 16 h have been reported in Per1/Per2-deficient liver (Aviram et al. 2021). For each method, we reported the fraction of challenge datasets in which the inferred phases agreed with collection times more closely than expected by chance, assessed using the one-sided MAE permutation test ($p < 0.05$).

##### 1.5.5 Gene-level rhythm reconstruction

To assess gene-level rhythm reconstruction, a cosinor model was fitted to each gene's $log_2(TPM + 1)$ expression values using the inferred sample phases and, separately, the reference times. For GSE108539, which contains repeated biopsies from 10 donors, both fits included donor-specific intercepts as fixed effects, while the sine and cosine coefficients were shared across donors.

For the reference-time fits, the sine and cosine terms were tested jointly using a partial F-test. Genes with Benjamini–Hochberg-adjusted q values below 0.05 met the significance criterion for reference rhythmicity. For the main gene-level analyses, the reference-rhythmic set was restricted to genes with an acrophase standard error of at most 1.0 h. Standard errors were estimated using the delta method with the debiased amplitude.

As a sensitivity analysis, the gene-level measures below were also computed with two other definitions of reference rhythmicity: the q-value criterion alone, and the q-value criterion combined with a minimum cosinor amplitude of 0.20 or 0.29 log2 units in place of the standard-error criterion. The higher floor corresponds to a peak-to-trough ratio of approximately 1.5 on the $TPM + 1$ scale ($2^{2\times0.29} \approx 1.495$).

Three quantities were compared:

- the median absolute acrophase error over the reference-rhythmic genes, after the one rotation and reflection that minimize that median over the same genes (searched over the whole circle, to within 0.025°);
- the Spearman correlation of reconstructed amplitude with reference amplitude, over the same reference-rhythmic genes, excluding genes whose debiased reference amplitude is zero;
- rhythmicity reconstruction, assessed using the area under the receiver operating characteristic curve (AUC), with each gene's $R^2$ from the inferred-phase fit as the score and its reference rhythmicity classification as the label. For GSE108539, the score was the partial $R^2$ of the rhythm terms relative to a model containing donor intercepts alone. Genes classified as rhythmic by the q-value criterion but excluded by the acrophase standard-error criterion (or, in the sensitivity analysis, by an amplitude floor) were omitted from the AUC calculation rather than counted as non-rhythmic.

For the example in Fig. 3d, the acrophase error was the mean circular distance over the twelve clock genes after the rotation and reflection that minimize that mean, which differs from the median-error objective above.

Each quantity was summarized per method by its mean and median over the datasets of the main accuracy comparison, with family-weighted values alongside.

Each metric was compared with its permutation distribution, generated using 200 permutations of the inferred sample phases. For GSE108539, phases were permuted within donors. Note that because the same expression matrix was used for phase inference and gene-level evaluation, these analyses assessed rhythm reconstruction and did not provide independent validation of rhythmicity detection.

The same permutation of inferred sample phases was applied to all genes, preserving relationships among their expression profiles. In strongly rhythmic datasets, the fitted gene acrophases could retain some of their relative ordering. After aligning the fitted and reference acrophases with a single rotation and optional reflection applied to all genes, acrophase errors could remain small even under permutation. In simulations, shuffled sample phases produced median acrophase errors of approximately 1.6 h with strong rhythms and 2.9 h with weak rhythms, compared with 5.5 h without rhythms. Acrophase errors were therefore interpreted relative to the permutation distribution for each dataset.

##### 1.5.6 Method agreement and consensus

Common frame without reference times. To compare or combine methods, each method's phases on a dataset were brought into a common frame without using reference times: every method was aligned (rotation and reflection minimizing the circular MAE between the two) to one reference method, the circular mean of the aligned phases became the new reference, and the alignment was repeated once. Each method was used at its predefined configuration, and a method without output on a dataset was omitted there.

Agreement. For each pair of methods on a dataset, agreement was the mean resultant length of the residuals of one method's phases against the other's, after the rotation and reflection that maximize it (1 for phases identical up to rotation and reflection; unrelated phases give small positive values that shrink with sample size). A dataset's agreement was the mean over its pairs. Agreement was related to the dataset's mean aligned MAE across datasets with a reference time, with the challenge datasets included and marked.

Consensus. A consensus phase was the circular mean of the aligned phases of the top three, the top five, or all methods. Which methods were "top" was decided with the evaluated dataset's whole study family held out: methods were ranked by their mean aligned circular MAE over the datasets of the other families, using available values. The consensus was scored by aligned circular MAE and compared with three single-method references: the method ranked first under the same held-out ranking, the median MAE of the methods that ran on the dataset, and the best method on that dataset, which uses its reference times and is shown only as an upper bound. The challenge datasets were excluded from this comparison.

##### 1.5.7 Datasets without collection times

For the Diversity Outbred cohort (468 adipose and 481 liver samples; 466 mice in both), a cosinor was fitted to each gene against the method's phases. Evaluation was restricted to genes that were present in the expression matrix and had reference acrophase estimates. Agreement with the reference acrophases was assessed using median absolute acrophase error and circular correlation. Acrophase error was minimized over a single rotation and optional reflection applied to all evaluated genes. Both measures were compared with their permutation distributions, generated using 200 permutations of the inferred sample phases and the same gene set for observed and permuted scores. The primary reference consisted of acrophases from the corresponding tissue in the Zhang atlas (Zhang et al. 2014). CYCLOPS, CYCLOPS 2.0, DCPR and TEMPO used seed gene lists or priors derived from this atlas. Because these methods used information derived from the Zhang atlas, their agreement with its reference acrophases should not be interpreted as independent validation.

For liver, an additional set of reference acrophases was estimated from the mouse liver atlas (Brooks et al. 2023) after excluding the Zhang liver study.

For the 466 mice profiled in both tissues, the absolute Fisher-Lee circular correlation was calculated between inferred liver and adipose phases. This comparison assessed consistency across tissues without requiring reference collection times.

#### 1.6 Sensitivity analyses

##### 1.6.1 Normalization and log transform

Each method was evaluated using all input matrices compatible with its model assumptions on 6 datasets chosen to span species, tissues and rhythm strengths: GSE151565, GSE201207_heart, GSE151565_Cortex, GSE202537_Caudate, simulated liver, simulated hypothalamus. Log-transformed inputs for CYCLOPS and CYCLOPS 2.0, non-count inputs for TEMPO, and inputs without log transformation for CHIRAL and reCAT were classified as incompatible inputs. The CYCLOPS and CYCLOPS 2.0 runs with log-transformed inputs were reported but excluded from input selection. The incompatible TEMPO, CHIRAL and reCAT configurations were not run. To estimate what choosing the input by score would gain, each method's input was chosen by mean error with the held-out dataset's whole study family removed, then scored on the held-out dataset, and compared with the predefined configuration on the same datasets.

##### 1.6.2 Random seed

Each method included in the main rankings was rerun with 16 random seeds, beginning with the seed used in the main comparison. Each method was run using its predefined configuration on 7 datasets chosen to span species, tissues and rhythm strengths: GSE151565, GSE151565_Cortex, GSE201207_heart, GSE202537_Caudate, simulated hypothalamus, simulated liver, simulated liver with 24 samples. For each method and dataset, the standard deviation and range of aligned circular MAE were calculated across seeds. The median of each measure was then reported across the 7 datasets.

##### 1.6.3 Seed gene list

CHIRAL, CYCLOPS, CYCLOPS 2.0, DCPR and TEMPO were evaluated using three seed gene lists: the default list used by each method in the main comparison, the list of 3,186 rhythmic liver genes from the Zhang atlas, and the twelve clock genes. This analysis used the datasets described in the random-seed analysis, excluding the simulation with 24 samples. For each method, comparisons were restricted to datasets with evaluable outputs for all three lists. For CYCLOPS and CYCLOPS 2.0 the method's default list is the tissue-matched list wherever the main comparison uses one. On the liver datasets the tissue list of CYCLOPS is the Zhang liver list itself, so for CYCLOPS its default list and the Zhang liver list are the same list there, and that comparison is identical by construction (for CYCLOPS 2.0 the default list is the union of the tissue list with the ubiquity list, so the two differ). Follow-up: decide how to treat this comparison. The 6 datasets belonged to four study families. For each method, paired differences in aligned circular MAE between each seed gene list and the method's default list were summarized by their mean and median with equal dataset weights, and by family-weighted mean and median, giving each study family equal total weight. These differences were reported descriptively; no hypothesis tests were performed.

##### 1.6.4 Tissue-specific seed gene lists

To assess tissue-specific seeding, CYCLOPS and CYCLOPS 2.0 were also run using their respective ubiquity seed gene lists alone on datasets with a corresponding tissue in the Zhang atlas. Within each method, runs with and without tissue-specific seeding differed only in the seed gene list. The atlas-derived tissue lists contained more genes than the approximately 35 genes in the default tissue-specific component of CYCLOPS 2.0. Because the atlas did not provide gene-level p values for these lists, they could not be restricted using a p-value threshold.

##### 1.6.5 Batch and covariate adjustment

In the main comparison, methods received expression data without sample-level batch or covariate information. We evaluated three approaches to batch and covariate adjustment in separate sensitivity analyses using datasets with the relevant annotations (Methods Table 5). Published workflows differed in their treatment of covariates. CHIRAL removed covariate-associated variation before phase inference (Talamanca et al. 2023), whereas CYCLOPS 2.0 included covariates in its model and its authors recommended against removing their effects before inference (Li et al. 2024). Covariate adjustment was not used in the published analyses of COFE, PENN and PROTECT (Ananthasubramaniam and Venkataramanan 2025; Ansary Ogholbake and Cheng 2024, 2025). The remaining methods did not describe a covariate-adjustment procedure.

ComBat-seq adjustment. Counts were adjusted using ComBat-seq (Zhang et al. 2020), with one annotation designated as the batch variable and the other covariates listed in Methods Table 5 entered as additional model terms. No biological group was specified for preservation during adjustment. Normalized input matrices were then recalculated from the adjusted counts, and each method was run using its predefined configuration.

Covariate adjustment within CYCLOPS 2.0. CYCLOPS 2.0 received the categorical covariates listed in Methods Table 5, which entered its model as separately encoded terms (Li et al. 2024). Before training, eigengenes were also regressed on the first listed covariate, and those largely explained by it were downweighted. This step accepted only one covariate. Because the authors did not specify which covariate to use, we placed the ComBat-seq batch variable first. Continuous covariates were not included because CYCLOPS 2.0 did not support their adjustment.

Covariate regression before CHIRAL. Following CHIRAL's published workflow, expression values for each gene were regressed on the covariates using ordinary least squares, and the residuals were used as input to CHIRAL. This procedure was applied only to CHIRAL. CHIRAL's own workflow regresses four covariates (ischaemic time, age, sex and death type); here the regression uses the same variables as the other two approaches, so that differences between approaches reflect the adjustment method rather than the choice of variables.

Methods Table 5. Variables supplied to each batch and covariate adjustment approach for each dataset. An asterisk identifies the covariate used in CYCLOPS 2.0's eigengene regression. Age group and disease state were biological annotations, post-mortem interval described tissue handling, and batch denoted the sequencing run in the Diversity Outbred cohort or the introduced batch effect in simulations. "Not applied" indicates that the adjustment approach was not used.

| Dataset | ComBat-seq batch | ComBat-seq additional covariates | CYCLOPS 2.0 covariates | CHIRAL regression |
|---|---|---|---|---|
| GSE202537_Caudate | Not applied | Not applied | Not applied | post-mortem interval, binned |
| GSE202537_Caudate_all | disease state (control / psychosis) | post-mortem interval, binned | disease state (control / psychosis)*, post-mortem interval, binned | disease state (control / psychosis), post-mortem interval, binned |
| GSE202537_Putamen | Not applied | Not applied | Not applied | post-mortem interval, binned |
| GSE202537_Putamen_all | disease state (control / psychosis) | post-mortem interval, binned | disease state (control / psychosis)*, post-mortem interval, binned | disease state (control / psychosis), post-mortem interval, binned |
| GSE201207_adrenal | age group (3 levels) | none | age group (3 levels)* | age group (3 levels) |
| GSE201207_heart | age group (3 levels) | none | age group (3 levels)* | age group (3 levels) |
| GSE201207_hypothalamus | age group (3 levels) | none | age group (3 levels)* | age group (3 levels) |
| GSE201207_kidney | age group (3 levels) | none | age group (3 levels)* | age group (3 levels) |
| GSE201207_lung | age group (3 levels) | none | age group (3 levels)* | age group (3 levels) |
| GSE201207_muscle | age group (3 levels) | none | age group (3 levels)* | age group (3 levels) |
| sim liver-balanced | batch | none | batch* | batch |
| sim liver-mild | batch | none | batch* | batch |
| GSE266549_adipose | batch | sex, generation | batch*, sex, generation | sex, generation, batch |
| GSE266569_liver | batch | sex, generation | batch*, sex, generation | sex, generation, batch |

Each adjusted run is paired with the unadjusted run of the same method, dataset and configuration. Where adjustment approaches were compared, the same covariates were supplied, but each pipeline incorporated them differently: ComBat-seq adjusts counts, CHIRAL receives regression residuals, and CYCLOPS 2.0 includes covariates in its own model. A difference between approaches is therefore an effect of the adjustment pipeline as implemented. None of the adjustment approaches knows the true phase, so an adjustment can remove circadian variation whenever a covariate is associated with collection time. We measured that association for each covariate as the fraction of the variance of $(cos\theta, sin\theta)$ explained by a linear model on it, $1 - \sum\|r_i\|^2/\sum\|y_i - \bar{y}\|^2$ summed over both components, with categorical covariates dummy-coded. These values were reported, and were also used once in a design decision: when the covariate set for GSE202537 had to be reduced because ComBat-seq was numerically unstable with all five candidate covariates, binned post-mortem interval was retained as the only covariate with a substantial association with time of death. That choice therefore drew on the reference times.

### Supplementary Figures

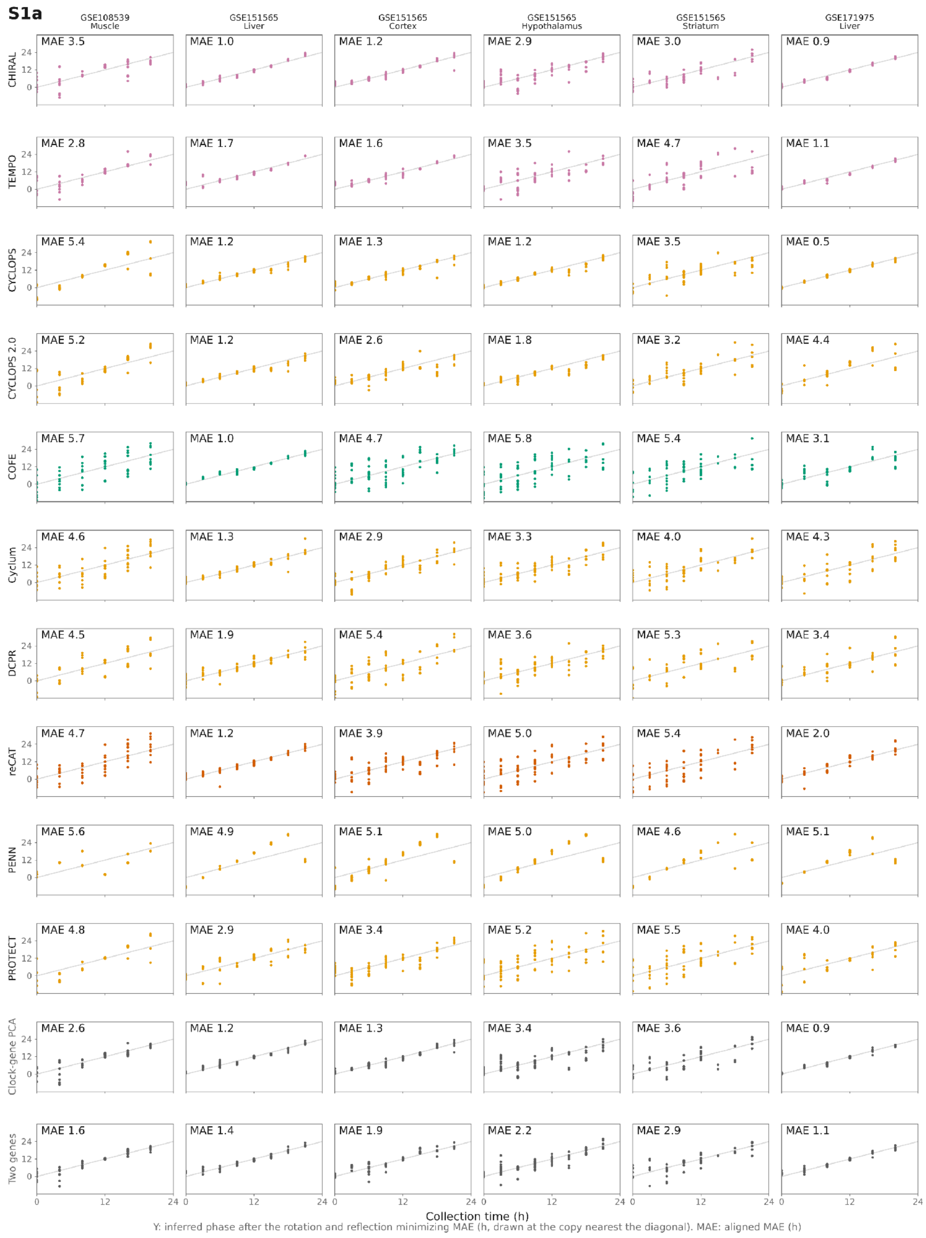


Figure S1a. Inferred phase against collection time, datasets of GSE108539, GSE151565 and GSE171975. Rows show methods and columns show datasets. Inferred phases were aligned to collection times by rotation and reflection to minimize circular MAE. Equivalent phase values differing by 24 h were plotted nearest the diagonal, which indicates agreement after alignment. MAE in each panel is the aligned MAE in hours.

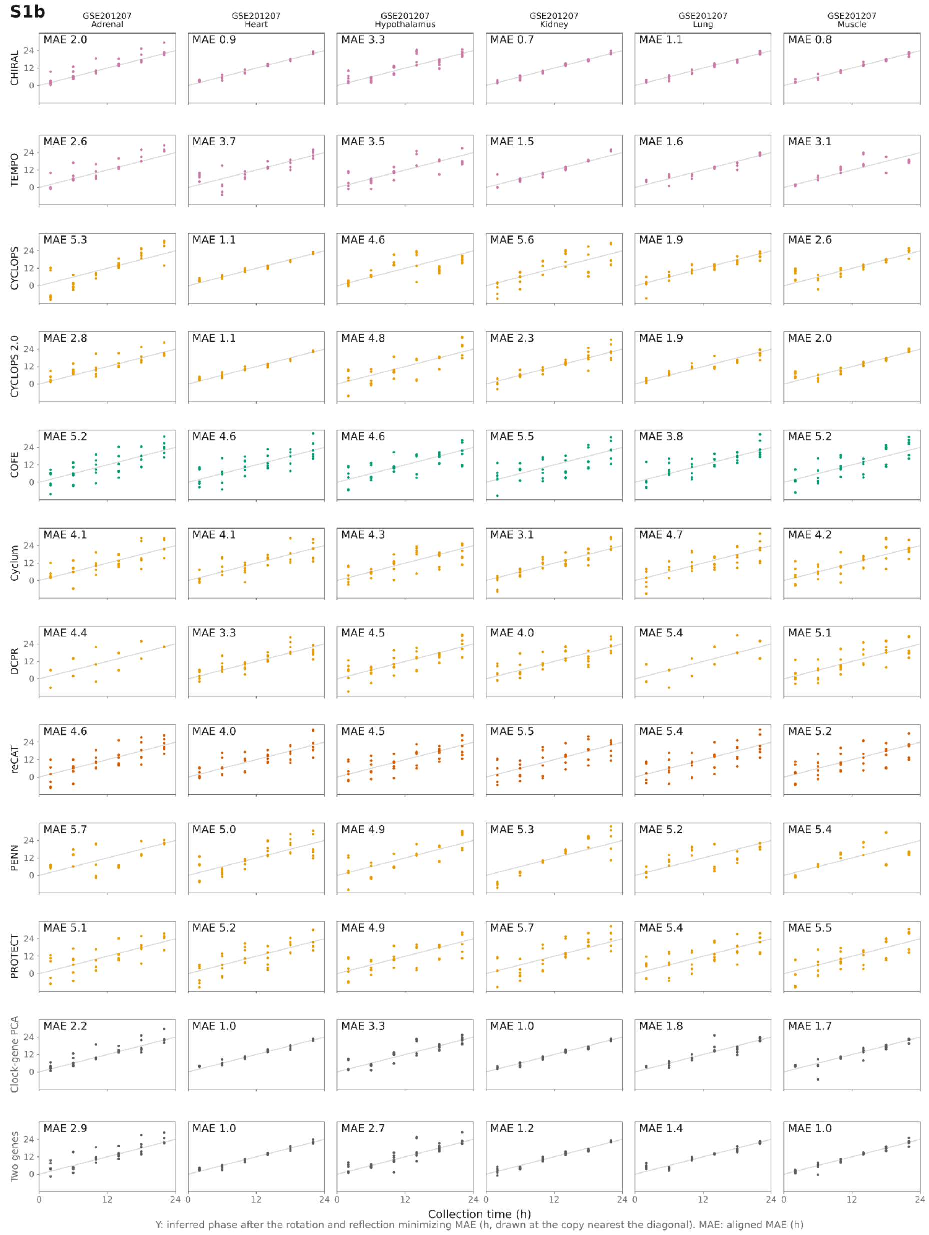


Figure S1b. Inferred phase against collection time, datasets of GSE201207. Rows show methods and columns show datasets. Inferred phases were aligned to collection times by rotation and reflection to minimize circular MAE. Equivalent phase values differing by 24 h were plotted nearest the diagonal, which indicates agreement after alignment. MAE in each panel is the aligned MAE in hours.

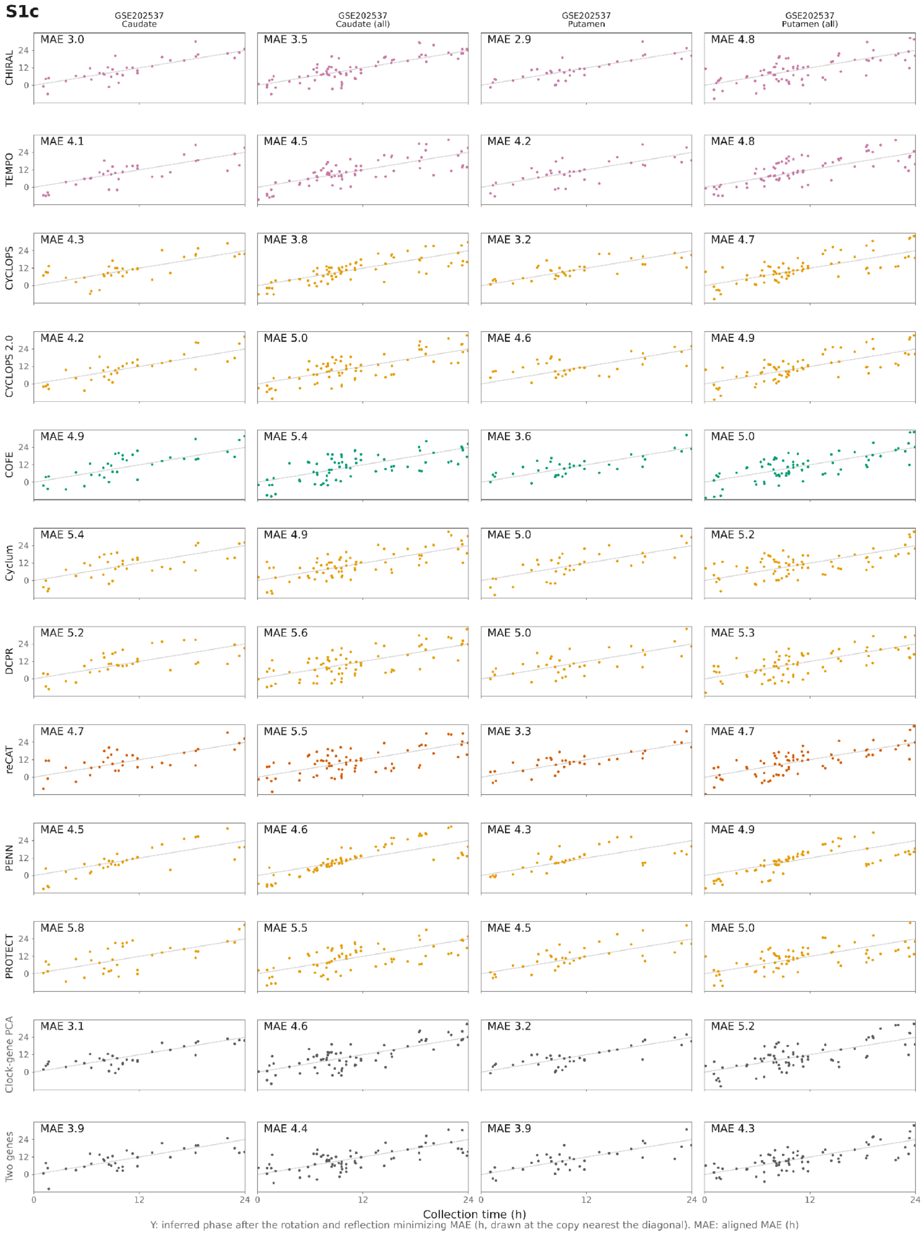


Figure S1c. Inferred phase against collection time, datasets of GSE202537. Rows show methods and columns show datasets. Inferred phases were aligned to collection times by rotation and reflection to minimize circular MAE. Equivalent phase values differing by 24 h were plotted nearest the diagonal, which indicates agreement after alignment. MAE in each panel is the aligned MAE in hours.

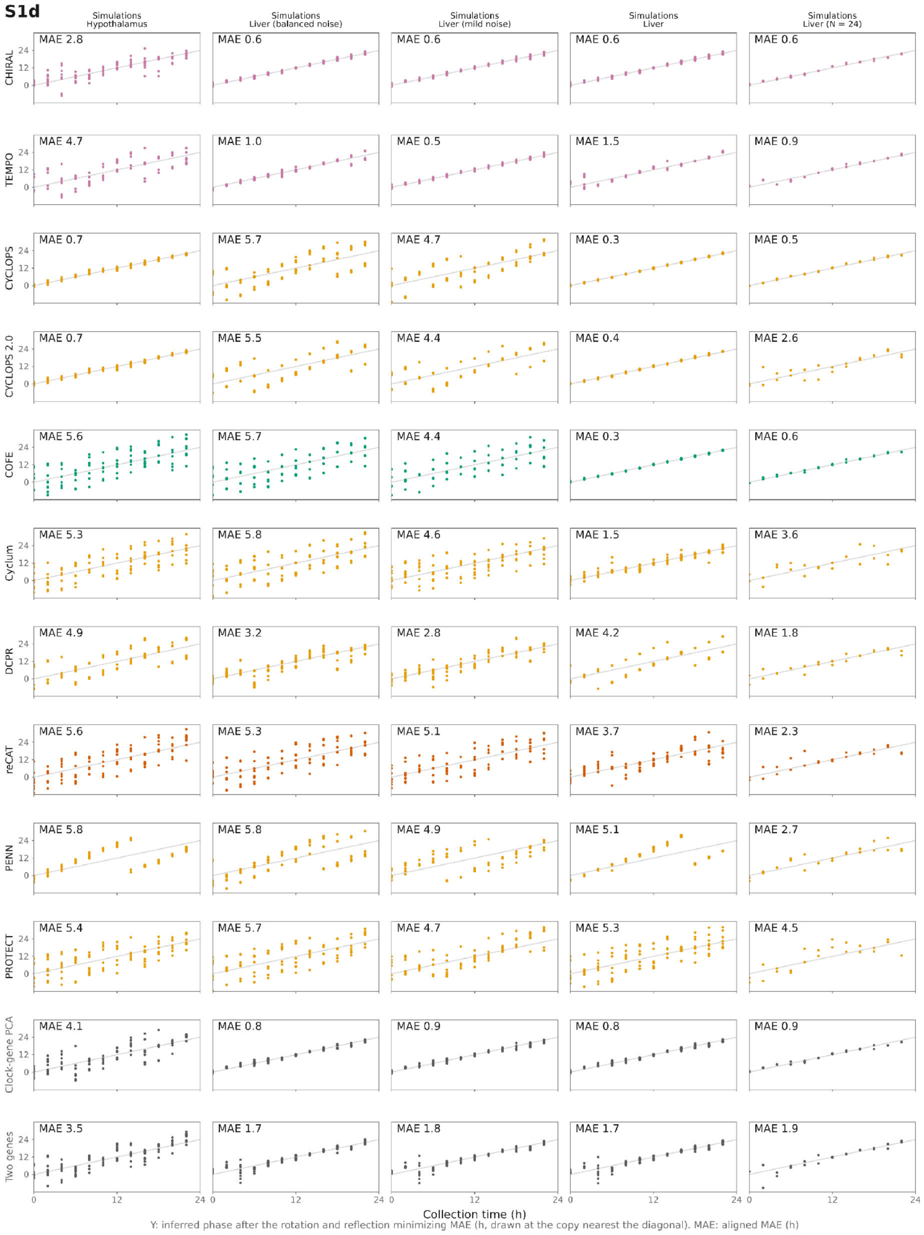


Figure S1d. Inferred phase against collection time, simulated datasets (1 of 2). Rows show methods and columns show datasets. Inferred phases were aligned to collection times by rotation and reflection to minimize circular MAE. Equivalent phase values differing by 24 h were plotted nearest the diagonal, which indicates agreement after alignment. MAE in each panel is the aligned MAE in hours.

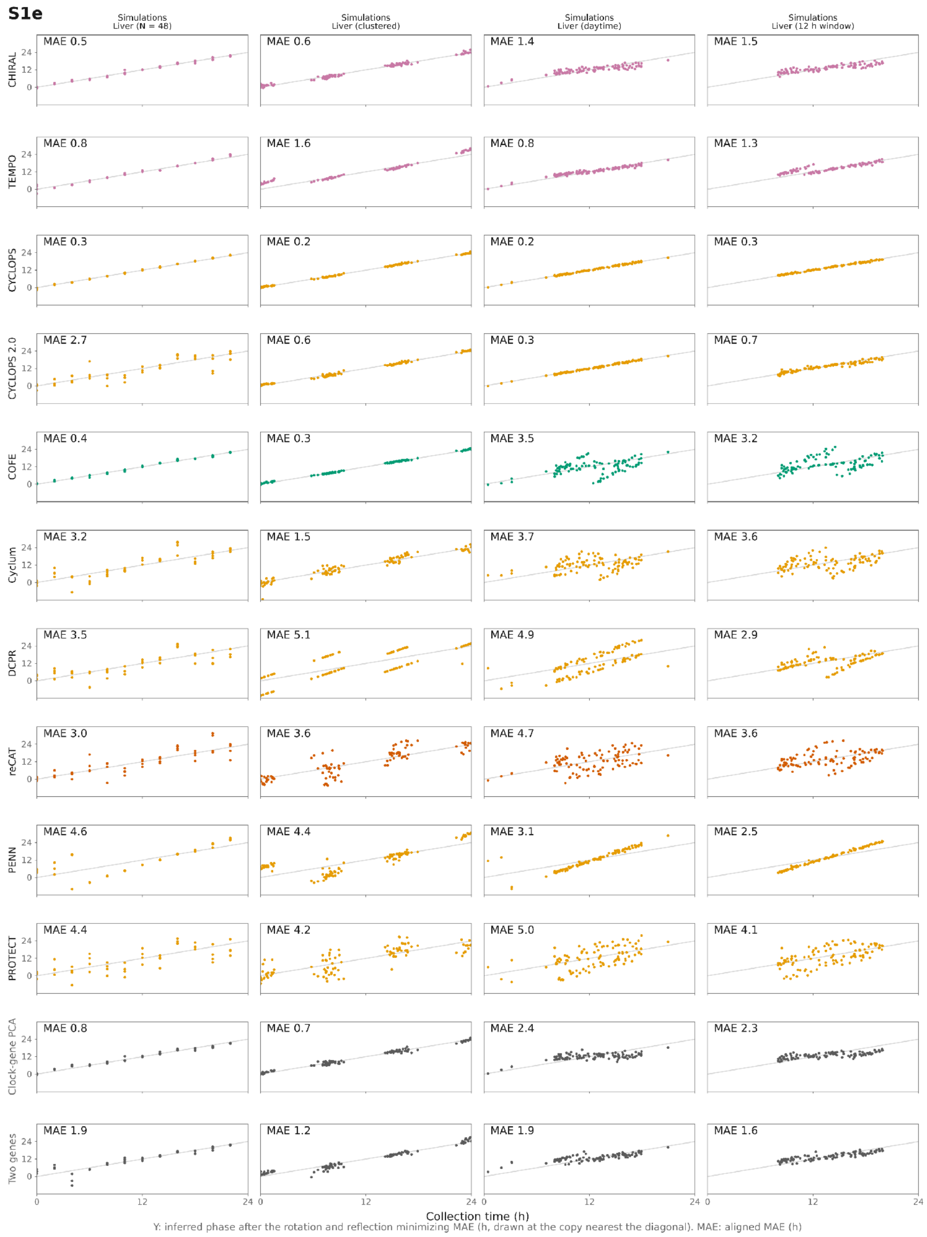


Figure S1e. Inferred phase against collection time, simulated datasets (2 of 2). Rows show methods and columns show datasets. Inferred phases were aligned to collection times by rotation and reflection to minimize circular MAE. Equivalent phase values differing by 24 h were plotted nearest the diagonal, which indicates agreement after alignment. MAE in each panel is the aligned MAE in hours.

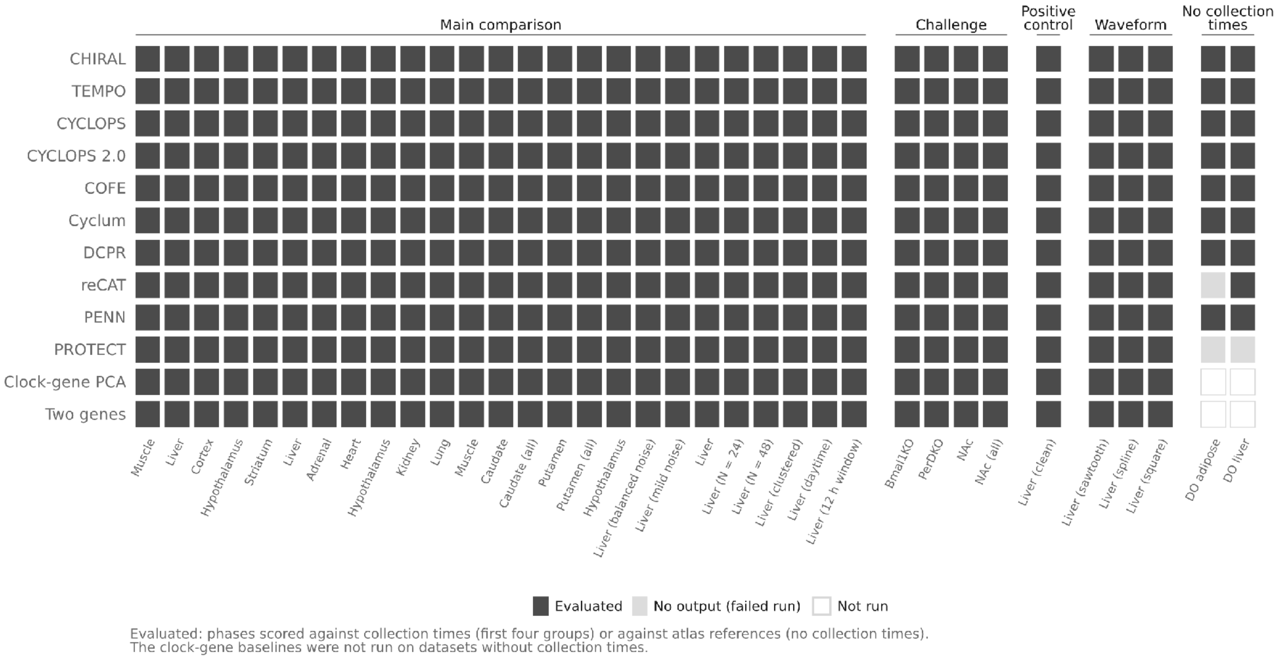


Figure S2. Availability of method outputs across datasets. For the 33 datasets with reference times, a run is evaluated when it returned phases for the labelled samples. No output was collapsed (at most 2 distinct phases; Fig. 2). For the 2 Diversity Outbred datasets, a run is evaluated when usable phases were available for the atlas comparison. No output means a failed run; every TEMPO run produced output, so none ended without a result after its prior sequence (Fig. S10). The clock-gene baselines were not run on datasets without collection times.

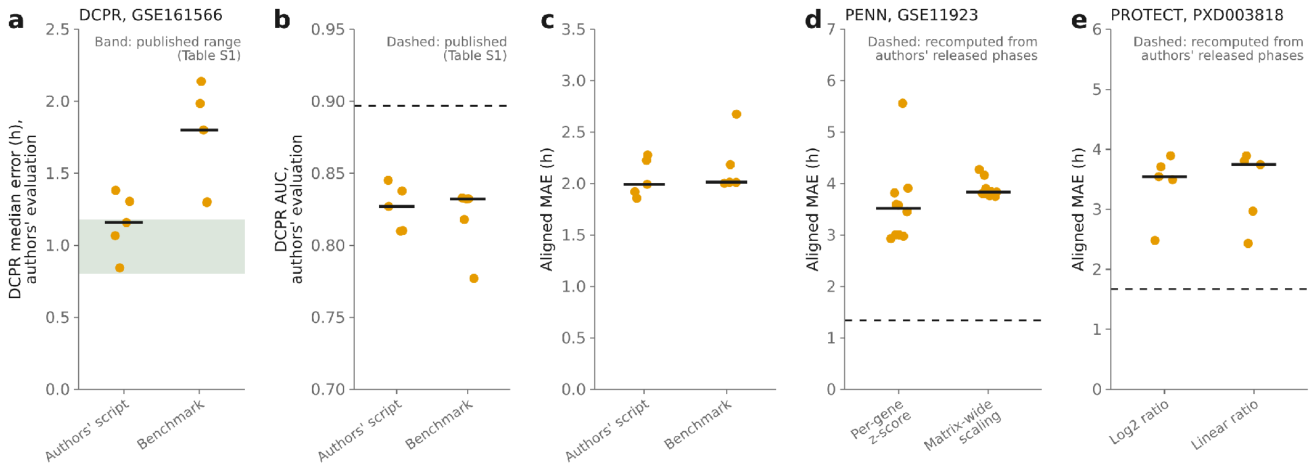


Figure S3. Comparisons with published results, one point per random seed (Table S1). (a–c) DCPR on GSE161566 with the authors' 528-gene list, run with the authors' script and with the benchmark implementation (5 and 5 seeds each; the authors' script has no seed setting and was run with the same seeds as the benchmark implementation): median error (a) and AUC (b) from the authors' evaluation code, with the published range (0.803–1.179 h, band) and value (0.897, dashed), sources in Table S1; (c) aligned MAE of the same runs, for which no published value exists. The authors' script aligns phases using the reference times before its evaluation; the benchmark uses the rotation and reflection minimizing MAE. (d) PENN on GSE11923 with the per-gene z-score used in the benchmark and with the code's matrix-wide scaling (10 seeds each). (e) PROTECT on PXD003818, reconstructed from the deposited protein table, on log2 and linear sample/reference ratios (5 seeds each). Dashed lines in (d, e): aligned MAE recomputed from the authors' released phase estimates and reference times. Bars: medians.

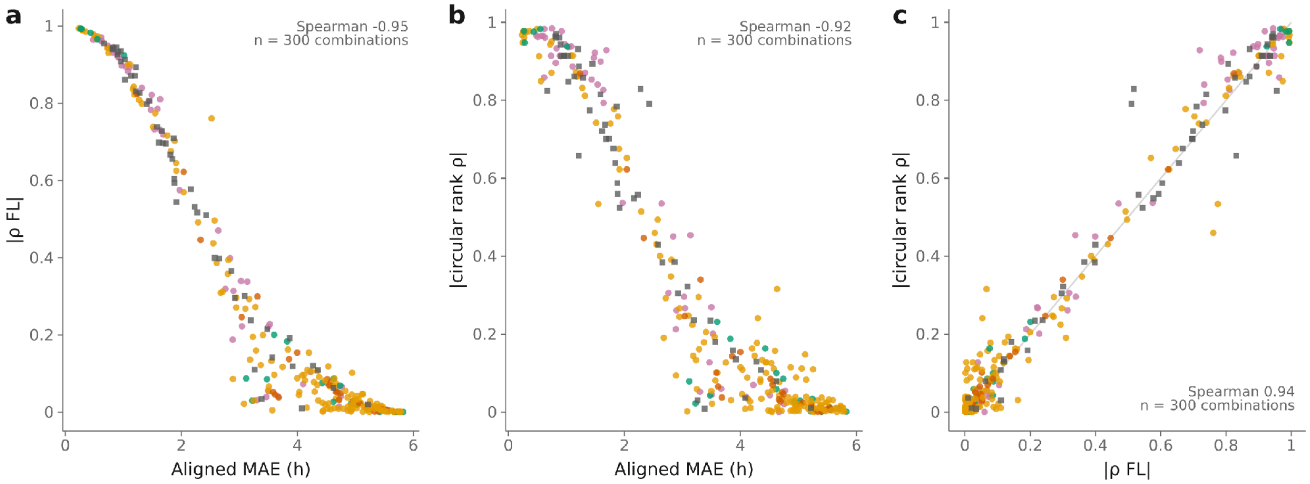


Figure S4. Agreement between the sample-level accuracy measures. Aligned MAE, absolute Fisher–Lee circular correlation and absolute circular rank correlation (which uses only the cyclic order of the samples, not their spacing), for each method-dataset combination of the main comparison (300 points: ten methods and two baselines on 25 datasets; colour: model family; squares: baselines): (a) MAE against Fisher–Lee correlation, (b) MAE against rank correlation, (c) Fisher–Lee against rank correlation, with Spearman correlations across combinations. The points share datasets and study families, so the correlations are descriptive.

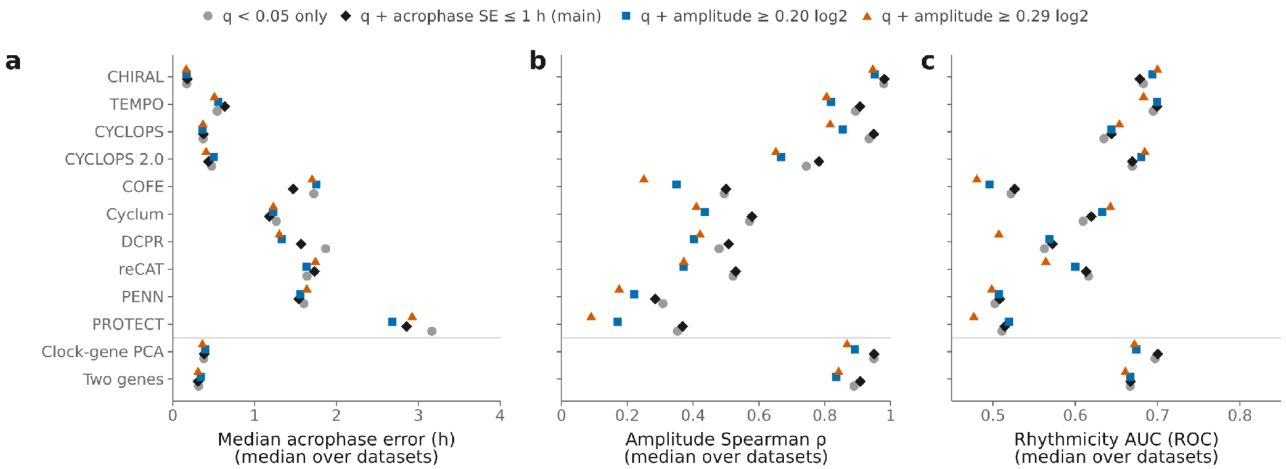


**d**

| Definition | Datasets with a defined acrophase error (of 25) | Reference-rhythmic genes, median over the 25 datasets | Range over datasets |
|---|---|---|---|
| q < 0.05 only | 22 | 1,900 | 0–9,289 |
| q + acrophase SE ≤ 1 h (main) | 22 | 1,517 | 0–6,051 |
| q + amplitude ≥ 0.20 log2 | 21 | 1,052 | 0–3,405 |
| q + amplitude ≥ 0.29 log2 | 21 | 386 | 0–1,742 |

Figure S5. Gene-level rhythm reconstruction under four definitions of reference rhythmicity. $q$ is the Benjamini–Hochberg-adjusted joint test of the cosine and sine terms in the cosinor fit to reference times. Definitions: $q$ < 0.05 alone; $q$ < 0.05 and acrophase standard error of at most 1.0 h (the main definition, Fig. 3); and $q$ < 0.05 and cosinor amplitude of at least 0.20 or 0.29 log2 units (on $log_2(TPM + 1)$). (a–c) Median over datasets of acrophase error, amplitude Spearman correlation and rhythmicity ROC AUC. (d) For each definition: the number of the 25 datasets in which at least one method has a defined acrophase error, and the median and range over all 25 datasets of the number of reference-rhythmic genes retained (datasets with none are included). The amplitude definitions leave one dataset fewer than the others. Restricting every definition to the 21 datasets scored under all four changes the plotted medians by at most 0.44 h for acrophase error, 0.04 for the amplitude correlation and 0.04 for the AUC.

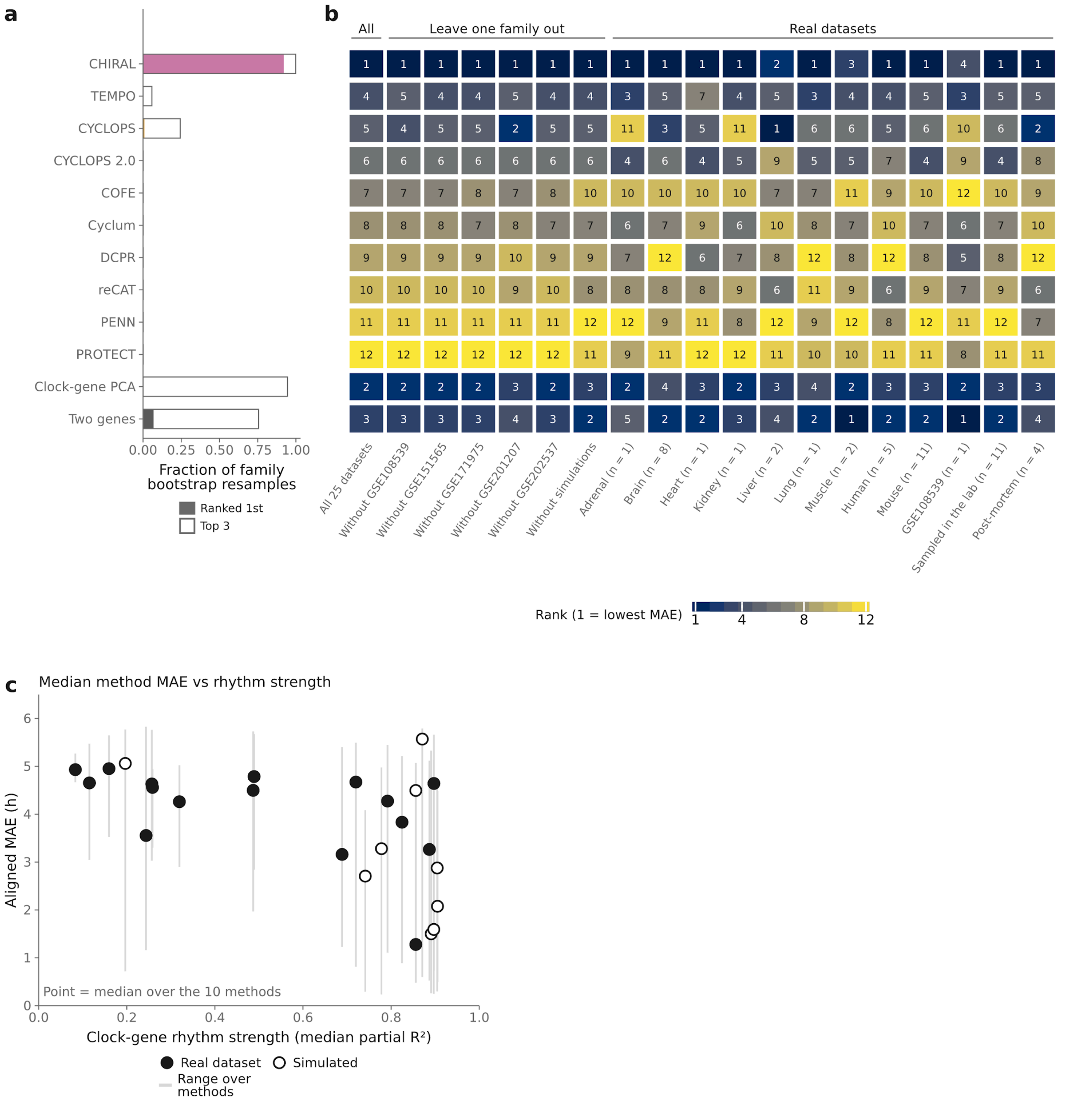


Figure S6. Sensitivity of method rankings to study composition. The ranking pool is the ten methods and two clock-gene baselines. (a) Fraction of 5,000 resamples of whole study families (6 families: five real and the simulations) in which each method had the lowest mean aligned MAE (filled) or ranked among the three lowest (outline). (b) Rank of each method by mean aligned MAE with equal dataset weights (1 = lowest, dark blue), recomputed on subsets of the main comparison: all 25 datasets; leaving out each of the 6 study families in turn; and, using only the 16 real datasets (“Real datasets”), by tissue, organism and source of reference times (sampled in the lab, post-mortem, and GSE108539, which was used in the development of CHIRAL and is its own study family, so leaving it out is also the analysis excluding GSE108539). n is the number of real datasets in a stratum; a stratum with one dataset ranks the methods on that dataset alone. The fractions describe the stability of the ranking within this benchmark, not the probability that a method is best in general. (c) Median aligned MAE over the 10 methods against clock-gene rhythm strength, one point per dataset of the main comparison (line: range over methods). Spearman correlation: -0.58 over all 25 datasets, -0.50 over the 16 real datasets and -0.52 over the 9 simulated datasets (descriptive; the median hides large differences between methods, shown by the range).

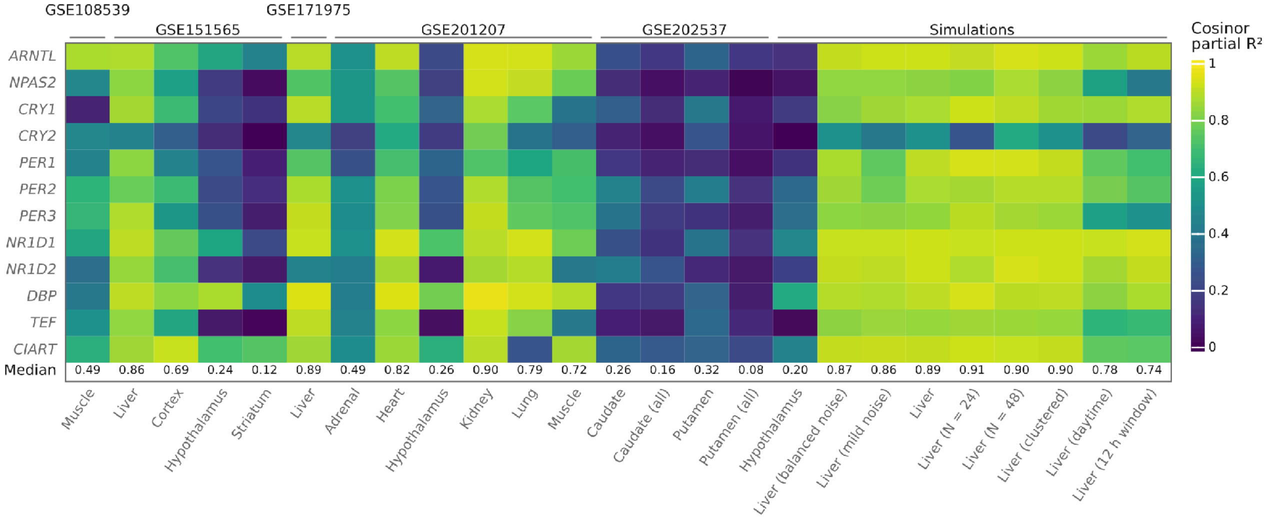


Figure S7. Clock-gene rhythm strength in the main-comparison datasets. Partial $R^2$ from cosinor fits of $log_2(TPM + 1)$ expression of each of twelve clock genes to collection time, with a separate donor intercept for GSE108539. The bottom row gives the median partial $R^2$ across the available clock genes, reported as the dataset's rhythm strength (Fig. 1b). –: gene absent from the dataset. Columns are grouped by study family.

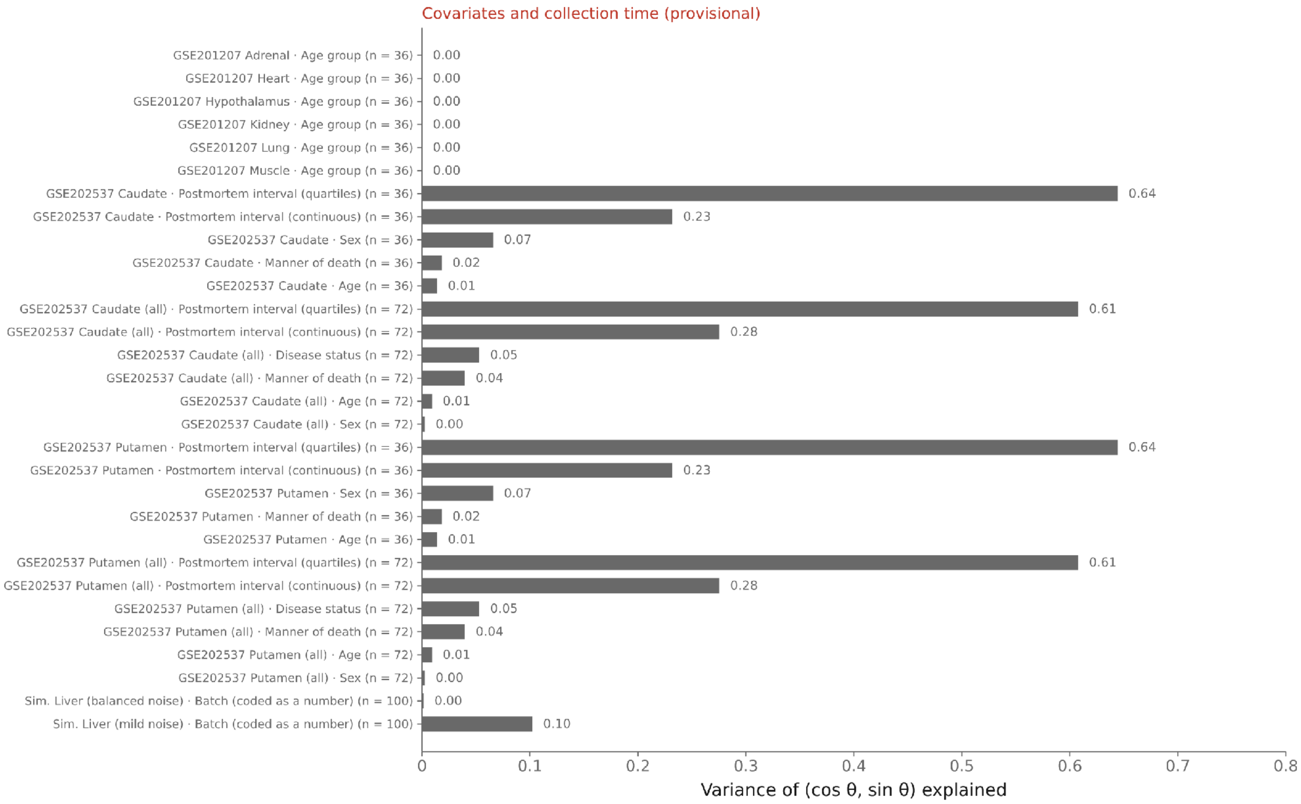


Figure S8. Association between covariates and collection times (provisional; red in the Methods). Proportion of the variance of $(cos\theta, sin\theta)$ explained by a separate linear model using each covariate as the predictor, where θ is the collection time expressed as an angle. The proportion was calculated from the combined residual and total sums of squares for the cosine and sine components (the unadjusted $R^2$, not corrected for the number of parameters); categorical covariates were dummy-coded and numeric covariates entered as one parameter. Postmortem interval is shown twice: continuous (one parameter) and in quartiles (equal-sized bins, three parameters), so the larger value for the binned version partly reflects its additional parameters. The batch covariate of the simulated datasets is coded as a number and fitted as one parameter. The caudate and putamen datasets come from the same donors, so their values are identical; n is the number of samples.

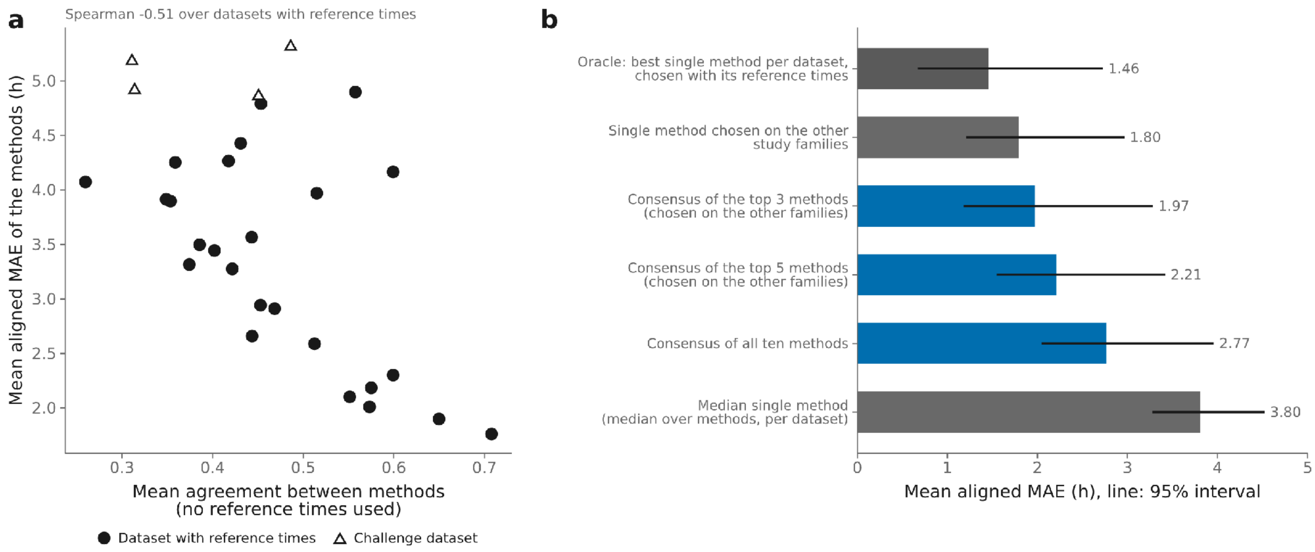


Figure S9. Between-method agreement and consensus inference. (a) Mean agreement between methods on each dataset, computed without reference times, against the methods' mean aligned MAE (triangles: challenge datasets). Agreement between two methods is the resultant length of the residuals after the best rotation and reflection of one onto the other, averaged over method pairs. (b) Mean aligned MAE over the 25 datasets of the main comparison (line: 95% interval from resampling whole study families) for: the consensus of the top 3, top 5 or all ten methods (circular mean of phases aligned to a common frame without reference times; clock-gene baselines excluded; the top 3 and top 5 chosen using the other study families only); the single method chosen using the other study families only; the median single method (the median over methods within each dataset, averaged over datasets); and, for comparison, the oracle, which selects the single method with the lowest MAE separately for each dataset using its reference times and so cannot be applied without them. The mean difference, consensus of the top 3 minus the single method chosen on the other families, is 0.17 h (95% interval -0.05 to 0.42 h, from resampling whole study families with the paired difference computed within each dataset; positive values favour the single method).

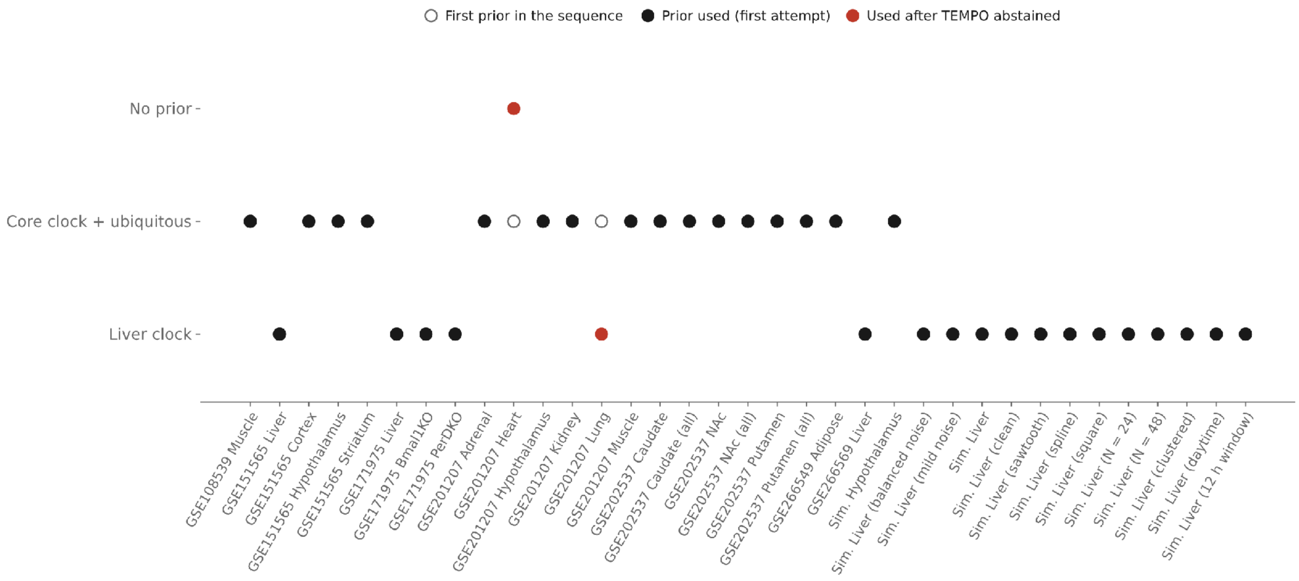


Figure S10. TEMPO prior sequence. For each dataset, the first prior in the sequence is shown as an open circle (a liver clock prior for liver datasets; otherwise a prior based on core clock genes and genes rhythmic across tissues). The filled circle marks the prior used in the successful run: black when it was the first attempt (the open circle is then hidden under it) and red when TEMPO abstained under earlier priors and the next prior in the sequence was used. TEMPO abstains when its fit is no better than shuffled phases; an abstention triggers the next prior, and any other error ends the sequence. Every run ended with a result, so no dataset reached the end of the sequence without one.

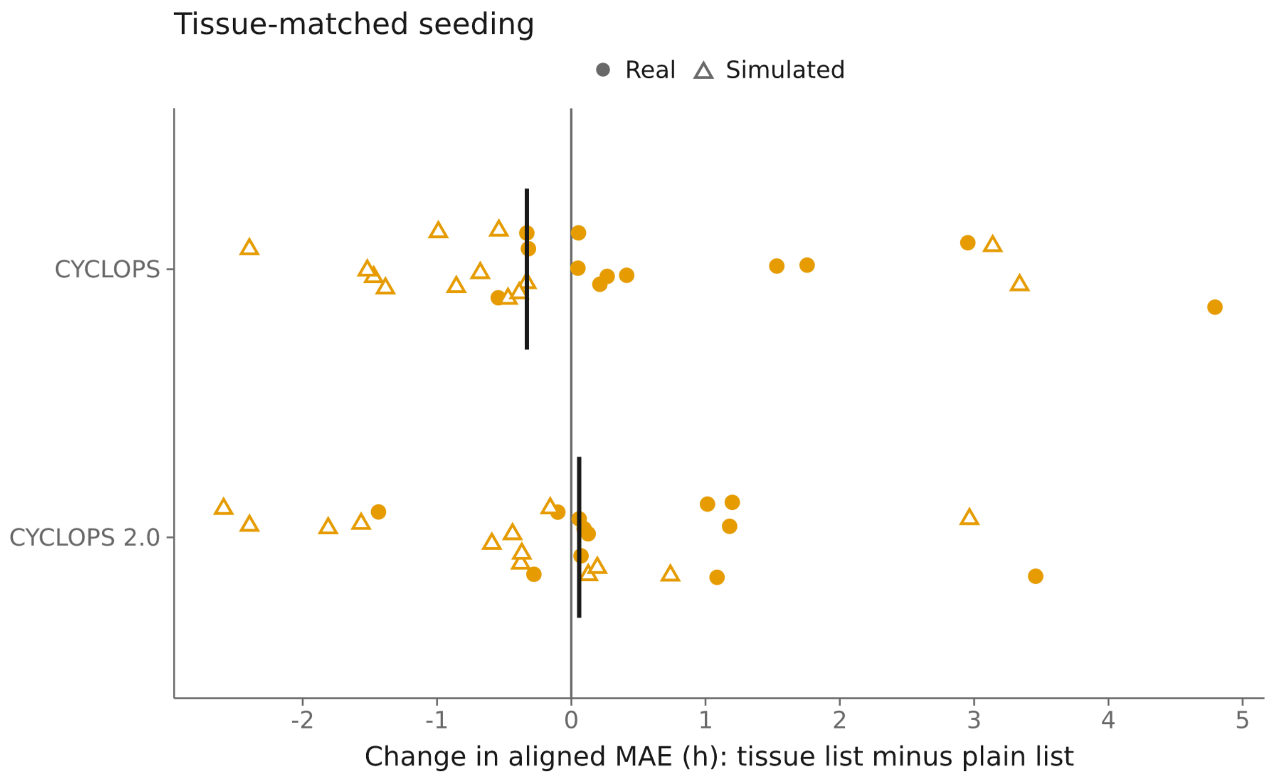


Figure S11. Effect of tissue-matched seed-gene lists. Change in aligned MAE for CYCLOPS and CYCLOPS 2.0 when a tissue-matched seed-gene list is used rather than the plain (ubiquity) list: tissue-matched list minus plain list, so negative values indicate lower aligned MAE with the tissue-matched list. Each point is a dataset for which a tissue-specific atlas seed list was available (25 datasets per method: 12 real and 13 simulated, including the challenge, positive-control and waveform datasets; circles: real datasets; open triangles: simulated datasets); bars: medians. The datasets differ from the six used for the seed-list comparison in Fig. 4h, which are not restricted to datasets with an atlas tissue.

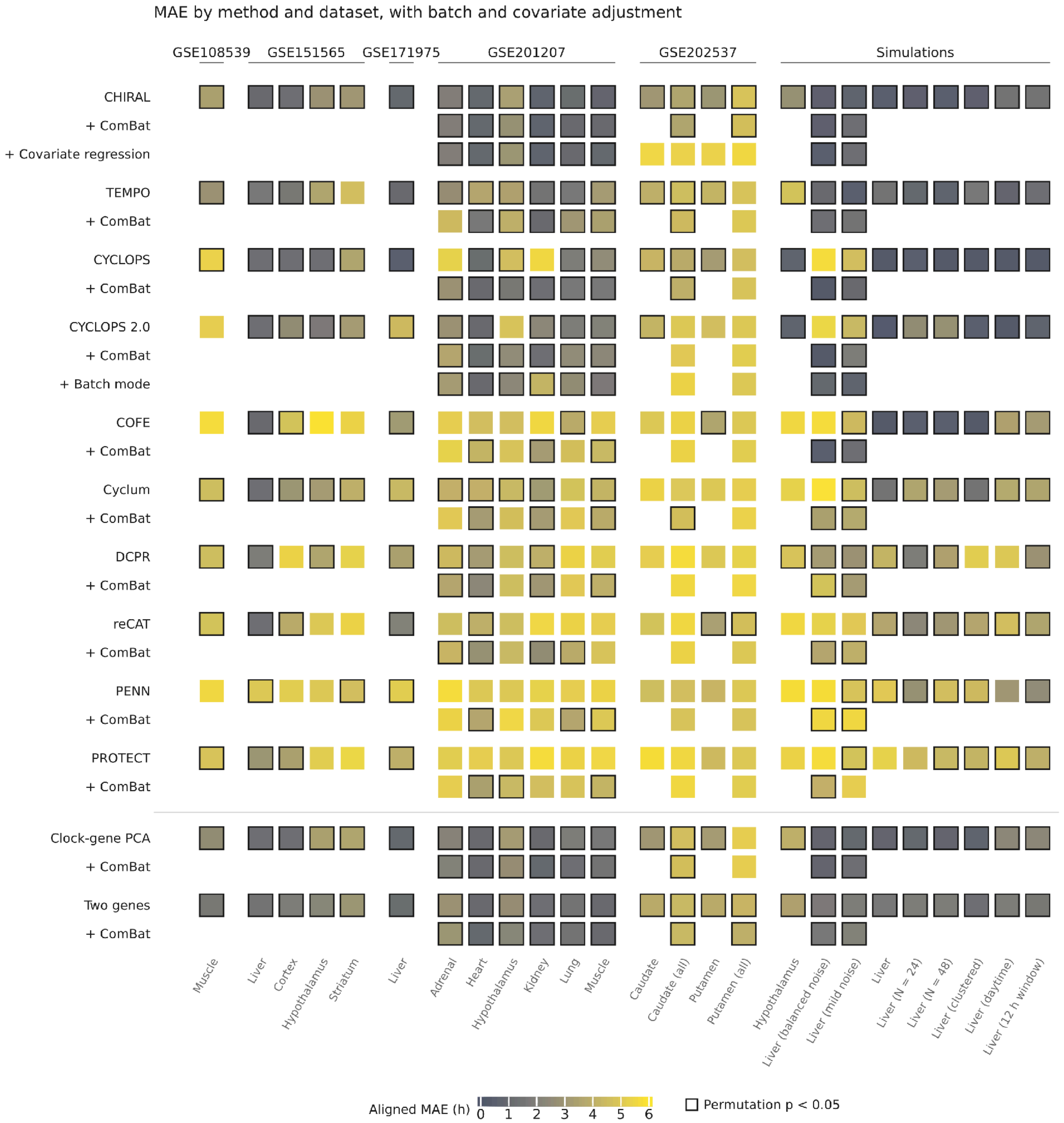


Figure S12. Aligned MAE by method and dataset, with batch and covariate adjustment. The heatmap of Fig. 2b with the adjusted runs added. Each method's unadjusted row is followed by the same method after ComBat-seq correction (10 datasets with batch labels) and, for CHIRAL, covariate regression (12 datasets) and, for CYCLOPS 2.0, its built-in batch mode (10 datasets). Blank cells indicate adjustments that were not evaluated. Fill: aligned MAE (h; lower is better). Dark outline: significant collection-time concordance (one-sided permutation $p < 0.05$, nominal).

#### Supplementary Tables

Table S1. Comparisons with published or released results. Methods were evaluated on the authors' datasets or example inputs before the main benchmark. The agreement criterion and any discrepancies are provided for each method. DCPR, PENN and PROTECT were evaluated with their final benchmark configurations at several random seeds, and their aligned errors use the exact MAE minimization described in Supplementary Methods, Aligned circular error. Aligned errors for the other methods used the earlier circular-mean rotation.

| Method | Reference result | Evaluation metric | Agreement criterion | Observed result | Comparison status |
|---|---|---|---|---|---|
| CYCLOPS | published median error, 1.69 h (BA11, probe level) | median absolute error, benchmark and authors' scripts | within the published value | 1.64 h with probe-level input and the rank-based seed rule | Published result reproduced using probe-level input and the published rank-based seed gene rule |
| CYCLOPS 2.0 | authors' released phases (BA11) | aligned error vs released phases | agreement after rotation and reflection | 0.00 h, R = 1.00 | Released phases matched after rotation and reflection |
| CHIRAL | authors' package call on their example | maximum difference in phases | numerical identity | $2 \times 10^{-14}$ | Numerical agreement with the authors' package; accuracy was not assessed |
| COFE | authors' notebook, 0.235 h | aligned error | equal to the notebook | 0.235 h | Notebook result reproduced after two corrections to the benchmark scripts |
| DCPR | published median error 0.803–1.179 h and AUC 0.897 (GSE161566, 12 samples, the authors' 528-gene list) | authors' evaluate (median error, AUC); aligned error | median error inside the published range | five seeds, median (range). Authors' script: median error 1.16 h (0.84–1.38), AUC 0.827 (0.810–0.845), aligned error 1.99 h. Benchmark scripts: median error 1.80 h (1.30–2.14), AUC 0.832 (0.777–0.833), aligned error 2.01 h | The authors' script reproduced the published median-error range on three of five seeds; the AUC was lower than published with both scripts. Scored with the same alignment, the two scripts gave similar aligned mean errors (1.99 and 2.01 h) and median errors of 1.49 and 1.80 h. The authors' script sets no random seed, so it was seeded here as the benchmark scripts are |
| PENN | authors' stored phases, 1.34 h against their labels (GSE11923, 48 | aligned error | none available | ten seeds, median (range): 3.52 h (2.93–5.56) with per-gene z-scoring; 3.83 h (3.75–4.27) | Not reproduced. Per-gene z-scoring did not account for the difference. Rerunning the |

| Method | Reference result | Evaluation metric | Agreement criterion | Observed result | Comparison status |
|---|---|---|---|---|---|
| | samples; no published error) | | | with the code's matrix-wide scaling | authors' notebook also did not reproduce their stored phases (3.21 h, earlier evaluation) |
| PROTECT | authors' stored phases, 1.67 h against their labels (PXD003818, 16 samples) | aligned error | none available | five seeds, median (range): 3.55 h (2.48–3.89) on log2 ratios; 3.75 h (2.43–3.89) on linear ratios | Not reproduced. The input was reconstructed from the deposited protein table (2,316 proteins quantified in all samples), because the authors' input file and filter are not published |
| TEMPO | authors' shipped fitted coefficients | maximum absolute difference | numerical tolerance | $5.6 \times 10^{-4}$ | Numerical agreement was assessed for fitted coefficients in the container; phase-inference accuracy was not assessed |
| Cyclum | authors' notebook precision, 0.679 | authors' precision estimate | close to the notebook | 0.690 (three dimensions) | Precision was close to the reported notebook value after a correction to the benchmark script |
| reCAT | authors' shipped ordering | aligned error, circular R | agreement | 1.10–1.12 h, R = 0.92 | Released ordering reproduced. The authors' multithreaded run was not deterministic |

Table S2. Summaries of every accuracy and gene-level measure for every method over the datasets of the main comparison: mean and median with equal dataset weights, family-weighted mean and median, means over real and simulated datasets, and the number of datasets with a value. Gene-level measures are first computed within each dataset.

| Measure | Method | Mean | Median | Family mean | Family median | Real mean | Simulated mean | n |
|---|---|---|---|---|---|---|---|---|
| aligned MAE (h) | CHIRAL | 1.8 | 1.23 | 2.08 | 1.54 | 2.23 | 1.03 | 25 |
| | TEMPO | 2.49 | 1.69 | 2.55 | 2.84 | 3.06 | 1.46 | 25 |
| | CYCLOPS | 2.52 | 1.91 | 2.77 | 2.55 | 3.13 | 1.44 | 25 |
| | CYCLOPS 2.0 | 2.8 | 2.61 | 3.5 | 4.45 | 3.26 | 1.98 | 25 |
| | COFE | 3.91 | 4.62 | 4.21 | 4.74 | 4.61 | 2.66 | 25 |
| | Cyclum | 3.94 | 4.13 | 4.11 | 4.35 | 4.1 | 3.64 | 25 |
| | DCPR | 4.22 | 4.52 | 4.24 | 4.54 | 4.51 | 3.7 | 25 |
| | reCAT | 4.26 | 4.66 | 4.03 | 4.69 | 4.36 | 4.1 | 25 |
| | PENN | 4.77 | 4.95 | 4.97 | 5.12 | 5.02 | 4.31 | 25 |
| | PROTECT | 4.86 | 5.01 | 4.73 | 4.83 | 4.9 | 4.79 | 25 |
| | Clock-gene PCA | 2.15 | 1.85 | 2.2 | 2.22 | 2.5 | 1.52 | 25 |
| | Two genes | 2.2 | 1.87 | 2.1 | 1.61 | 2.36 | 1.91 | 25 |
| | rho_FL | | CHIRAL | 0.64 | 0.83 | 0.55 | 0.73 | 0.52 |
| | TEMPO | 0.52 | 0.72 | 0.5 | 0.4 | 0.36 | 0.8 | 25 |
| | CYCLOPS | 0.53 | 0.65 | 0.48 | 0.44 | 0.39 | 0.77 | 25 |
| | CYCLOPS 2.0 | 0.44 | 0.39 | 0.29 | 0.01 | 0.33 | 0.64 | 25 |
| | COFE | 0.23 | 0.03 | 0.15 | 0.03 | 0.1 | 0.46 | 25 |
| | Cyclum | 0.17 | 0.08 | 0.14 | 0.08 | 0.14 | 0.21 | 25 |
| | DCPR | 0.11 | 0.02 | 0.12 | 0.1 | 0.08 | 0.16 | 25 |
| | reCAT | 0.13 | 0.05 | 0.2 | 0.05 | 0.15 | 0.1 | 25 |
| | PENN | 0.07 | 0.03 | 0.05 | 0.03 | 0.03 | 0.14 | 25 |
| | PROTECT | 0.04 | 0.01 | 0.05 | 0.03 | 0.04 | 0.02 | 25 |
| | Clock-gene PCA | 0.58 | 0.66 | 0.56 | 0.55 | 0.48 | 0.74 | 25 |
| | Two genes | 0.57 | 0.67 | 0.6 | 0.7 | 0.54 | 0.63 | 25 |
| | circular rank rho | | CHIRAL | 0.64 | 0.86 | 0.56 | 0.84 | 0.52 |
| | TEMPO | 0.56 | 0.79 | 0.54 | 0.45 | 0.4 | 0.83 | 25 |
| | CYCLOPS | 0.52 | 0.68 | 0.47 | 0.43 | 0.39 | 0.76 | 25 |
| | CYCLOPS 2.0 | 0.45 | 0.4 | 0.31 | 0.13 | 0.36 | 0.62 | 25 |
| | COFE | 0.24 | 0.04 | 0.16 | 0.03 | 0.11 | 0.46 | 25 |
| | Cyclum | 0.17 | 0.08 | 0.14 | 0.08 | 0.15 | 0.2 | 25 |
| | DCPR | 0.12 | 0.05 | 0.13 | 0.1 | 0.09 | 0.18 | 25 |
| | reCAT | 0.14 | 0.05 | 0.2 | 0.05 | 0.15 | 0.11 | 25 |
| | PENN | 0.08 | 0.02 | 0.07 | 0.01 | 0.05 | 0.15 | 25 |
| | PROTECT | 0.04 | 0.02 | 0.05 | 0.03 | 0.05 | 0.01 | 25 |

| Measure | Method | Mean | Median | Family mean | Family median | Real mean | Simulated mean | n |
|---|---|---|---|---|---|---|---|---|
| | Clock-gene PCA | 0.6 | 0.79 | 0.57 | 0.64 | 0.48 | 0.81 | 25 |
| | Two genes | 0.57 | 0.66 | 0.6 | 0.72 | 0.54 | 0.63 | 25 |
| median acrophase error (h) | CHIRAL | 0.54 | 0.18 | 0.88 | 0.28 | 0.74 | 0.26 | 22 |
| | TEMPO | 0.78 | 0.64 | 1.01 | 0.89 | 0.89 | 0.62 | 22 |
| | CYCLOPS | 0.52 | 0.38 | 0.76 | 0.41 | 0.7 | 0.25 | 22 |
| | CYCLOPS 2.0 | 1.2 | 0.44 | 1.33 | 0.58 | 1.04 | 1.43 | 22 |
| | COFE | 2.44 | 1.47 | 3.01 | 1.45 | 3.12 | 1.45 | 22 |
| | Cyclum | 1.47 | 1.18 | 1.47 | 1.2 | 1.22 | 1.84 | 22 |
| | DCPR | 2.29 | 1.57 | 2.21 | 0.96 | 2.43 | 2.11 | 22 |
| | reCAT | 1.93 | 1.73 | 1.76 | 1.83 | 1.88 | 2.01 | 22 |
| | PENN | 1.91 | 1.54 | 1.78 | 1.34 | 1.88 | 1.93 | 22 |
| | PROTECT | 3.36 | 2.86 | 3.39 | 2.45 | 3.83 | 2.67 | 22 |
| | Clock-gene PCA | 0.6 | 0.38 | 0.86 | 0.6 | 0.76 | 0.38 | 22 |
| | Two genes | 0.56 | 0.31 | 0.66 | 1 | 0.65 | 0.42 | 22 |
| amplitude Spearman rho | CHIRAL | 0.9 | 0.98 | 0.87 | 0.91 | 0.89 | 0.91 | 22 |
| | TEMPO | 0.81 | 0.91 | 0.79 | 0.9 | 0.78 | 0.85 | 22 |
| | CYCLOPS | 0.74 | 0.95 | 0.69 | 0.72 | 0.68 | 0.83 | 22 |
| | CYCLOPS 2.0 | 0.77 | 0.78 | 0.71 | 0.69 | 0.76 | 0.77 | 22 |
| | COFE | 0.52 | 0.5 | 0.49 | 0.58 | 0.41 | 0.67 | 22 |
| | Cyclum | 0.57 | 0.58 | 0.52 | 0.52 | 0.57 | 0.56 | 22 |
| | DCPR | 0.5 | 0.51 | 0.53 | 0.59 | 0.51 | 0.48 | 22 |
| | reCAT | 0.53 | 0.53 | 0.62 | 0.59 | 0.55 | 0.51 | 22 |
| | PENN | 0.35 | 0.28 | 0.41 | 0.39 | 0.31 | 0.39 | 21 |
| | PROTECT | 0.36 | 0.37 | 0.46 | 0.44 | 0.41 | 0.3 | 22 |
| | Clock-gene PCA | 0.87 | 0.95 | 0.86 | 0.88 | 0.86 | 0.88 | 22 |
| | Two genes | 0.84 | 0.91 | 0.77 | 0.9 | 0.82 | 0.87 | 22 |
| rhythmicity AUC (ROC) | CHIRAL | 0.74 | 0.68 | 0.67 | 0.64 | 0.63 | 0.91 | 22 |
| | TEMPO | 0.72 | 0.7 | 0.65 | 0.62 | 0.59 | 0.91 | 22 |
| | CYCLOPS | 0.67 | 0.64 | 0.59 | 0.58 | 0.54 | 0.87 | 22 |
| | CYCLOPS 2.0 | 0.69 | 0.67 | 0.59 | 0.6 | 0.58 | 0.84 | 22 |
| | COFE | 0.56 | 0.53 | 0.49 | 0.43 | 0.39 | 0.79 | 22 |
| | Cyclum | 0.62 | 0.62 | 0.57 | 0.57 | 0.55 | 0.72 | 22 |
| | DCPR | 0.57 | 0.57 | 0.52 | 0.41 | 0.44 | 0.76 | 22 |
| | reCAT | 0.56 | 0.61 | 0.57 | 0.64 | 0.44 | 0.73 | 22 |
| | PENN | 0.54 | 0.51 | 0.51 | 0.46 | 0.41 | 0.72 | 22 |
| | PROTECT | 0.5 | 0.51 | 0.45 | 0.37 | 0.37 | 0.68 | 22 |
| | Clock-gene PCA | 0.73 | 0.7 | 0.68 | 0.63 | 0.62 | 0.9 | 22 |

| Measure | Method | Mean | Median | Family mean | Family median | Real mean | Simulated mean | n |
|---|---|---|---|---|---|---|---|---|
| | Two genes | 0.73 | 0.67 | 0.65 | 0.62 | 0.61 | 0.9 | 22 |

Proceedings of the National Academy of Sciences 121 (7): e2311854121. https://doi.org/10.1073/pnas.2311854121.

Liang, Shaoheng, Fang Wang, Jincheng Han, and Ken Chen. 2020. “Latent Periodic Process Inference from Single-Cell RNA-Seq Data.” Nature Communications 11: 1441. https://doi.org/10.1038/s41467-020-15295-9.

Liu, Zehua, Huazhe Lou, Kaikun Xie, et al. 2017. “Reconstructing Cell Cycle Pseudo Time-Series via Single-Cell Transcriptome Data.” Nature Communications 8: 22. https://doi.org/10.1038/s41467-017-00039-z.

Love, Michael I., Wolfgang Huber, and Simon Anders. 2014. “Moderated Estimation of Fold Change and Dispersion for RNA-seq Data with DESeq2.” Genome Biology 15 (12): 550. https://doi.org/10.1186/s13059-014-0550-8.

Perrin, Laurent et al. 2018. “Transcriptomic Analyses Reveal Rhythmic and CLOCK-Driven Pathways in Human Skeletal Muscle.” eLife 7: e34114. https://doi.org/10.7554/eLife.34114.

Robinson, Mark D., and Alicia Oshlack. 2010. “A Scaling Normalization Method for Differential Expression Analysis of RNA-Seq Data.” Genome Biology 11: R25. https://doi.org/10.1186/gb-2010-11-3-r25.

Talamanca, Lorenzo, Cédric Gobet, and Felix Naef. 2023. “Sex-Dimorphic and Age-Dependent Organization of 24-Hour Gene Expression Rhythms in Humans.” Science 379 (6631): 478–83. https://doi.org/10.1126/science.add0846.

Tyler, Anna L. et al. 2025. “Transcripts with High Distal Heritability Mediate Genetic Effects on Complex Metabolic Traits.” Nature Communications 16: 5507. https://doi.org/10.1038/s41467-025-61228-9.

Wolff, C. A. et al. 2023. “Defining the Age-Dependent and Tissue-Specific Circadian Transcriptome in Male Mice.” Cell Reports 42 (1): 111982. https://doi.org/10.1016/j.celrep.2022.111982.

Wu, Gang, Ron C. Anafi, Michael E. Hughes, Karl Kornacker, and John B. Hogenesch. 2016. “MetaCycle: An Integrated R Package to Evaluate Periodicity in Large Scale Data.” Bioinformatics 32 (21): 3351–53. https://doi.org/10.1093/bioinformatics/btw405.

Yang, Jianing, Gregory R. Grant, and Thomas G. Brooks. 2025. “Generating Correlated Data for Omics Simulation.” PLOS Computational Biology 21 (9): e1013392. https://doi.org/10.1371/journal.pcbi.1013392.

Zhang, R., N. F. Lahens, H. I. Ballance, M. E. Hughes, and J. B. Hogenesch. 2014. “A Circadian Gene Expression Atlas in Mammals: Implications for Biology and Medicine.” Proceedings of the National Academy of Sciences of the USA 111 (45): 16219–24. https://doi.org/10.1073/pnas.1408886111.

Zhang, Y., G. Parmigiani, and W. E. Johnson. 2020. “ComBat-seq: Batch Effect Adjustment for RNA-seq Count Data.” NAR Genomics and Bioinformatics 2 (3): lqaa078. https://doi.org/10.1093/nargab/lqaa078.
